# Beyond the limits of the unassigned protist microbiome: inferring large-scale spatio-temporal patterns of marine parasites

**DOI:** 10.1101/2022.07.24.501282

**Authors:** Iris Rizos, Pavla Debeljak, Thomas Finet, Dylan Klein, Sakina-Dorothée Ayata, Fabrice Not, Lucie Bittner

## Abstract

Marine protists are major components of the oceanic microbiome that remain largely unrepresented in culture collections and genomic reference databases. The exploration of this uncharted protist diversity in oceanic communities relies essentially on studying genetic markers from the environment as taxonomic barcodes. Here we report that across 6 large scale spatio-temporal planktonic surveys, half of the genetic barcodes remain taxonomically unassigned at the genus level, preventing a fine ecological understanding for numerous protist lineages. Among them, parasitic Syndiniales (Dinoflagellata) appear as the least described protist group. We have developed a computational workflow, integrating diverse 18S rDNA gene metabarcoding datasets, in order to infer large-scale ecological patterns at 100% similarity of the genetic marker, overcoming the limitation of taxonomic assignment. From a spatial perspective, we identified 2 171 unassigned clusters exclusively shared between the Tropical/Subtropical Ocean and the Mediterranean Sea among all Syndiniales orders and 25 ubiquitous clusters shared within all the studied marine regions. From a temporal perspective, over 3 time-series, we highlighted 38 unassigned clusters that follow rhythmic patterns of recurrence and are the best indicators of parasite community’s variation. These clusters withhold potential as ecosystem change indicators, mirroring their associated host community responses. Our results underline the importance of Syndiniales in structuring planktonic communities through space and time, raising questions regarding host-parasite association specificity and the trophic mode of persistent Syndiniales, while providing an innovative framework for prioritizing unassigned protist taxa for further description.

## Introduction

The advances in high-throughput sequencing technologies have provided a new perspective to microbial diversity at a global scale. Studying the DNA of environmental microbial communities (i.e. microbiome) allowed to overcome the limit of non-cultivability and provided access to an unprecedented large quantity of high resolution genetic information [1-4]. *In silico* downstream analysis of genetic big-data shed light on a new challenge in microbial ecology: exploring the unassigned microbiome [5,6]. From the point of view of environmental genomics, the unassigned microbiome encompasses all genetic sequences that cannot be annotated with referenced biological information as they have no match in databases at a functional [6-9] and/or taxonomic level [5, 10, 11]. Recent studies have pointed out that unassigned sequences contribute to 25-58% of microbial communities’ diversity observed across a variety of aquatic and soil ecosystems [4, 5, 12]. In the marine realm, large scale sequencing studies have revealed that the unassigned microbiome represents half of the functional diversity (including samples enriched in viruses, prokaryotes and protists) [13]. In terms of taxonomic diversity, the unassigned protist microbiome, defined as taxa with V9 regions of the 18S rDNA marker having a sequence similarity <80% with reference sequences, represents ∼30% at the supergroup level ([14].

In marine metabarcoding studies, Syndiniales (a clade of marine alveolates, MALVs [15]) represent an ubiquitous and hyperdiverse lineage of protistan endoparasites [16-18]. Syndiniales are distributed worldwide from tropical and temperate zones [14, 19] to both arctic and antarctic poles [20, 21]. Their unexpected contribution to protist community composition has been revealed by metabarcoding studies both in open sea and coastal environments, with Syndiniales being the third most abundant lineage of the circumglobal Tara Oceans expedition [14] and representing up to 11% of community’s abundance in fjordic-bays [21] and 28% at a North-Atlantic river estuary [22]. Accumulating observations and correlations of metabarcoding data support that Syndiniales are opportunistically infecting a wide spectrum of hosts, including other protists (dinoflagellates, ciliates, radiolarians) but also metazoans (e.g. crustaceans) [22-23]. Their wide abundance and distribution confers them global ecological importance for microbial food webs and biogeochemical cycling, by regulating host populations [22, 24, 25] and supplying the microbial loop with organic matter [26]. Yet, the great majority of Syndiniales remain uncultivable and show a high degree of divergence in genomic sequences [27]. A recent study in an estuary revealed the existence of at least 8 cryptic Syndiniales species, among which 6 could be differentiated by the V4 region of the 18S marker by an 100% sequence similarity threshold [17]. Moreover, their complex lifestyle, small size (< 20 µm) and lack of distinctive morphological features makes Syndiniales’ description a laborious process relying on designing specific probes for *in situ* hybridization [24, 25, 28]. Thus, Syndiniales diversity still remains a blackbox in protistology [22, 25, 29], rendering the ecological understanding of these widespread microorganisms below the order level presently beyond reach [17].

In this study, we explored marine planktonic protist communities at a wide spatio-temporal scale, in order to: (i) quantify the taxonomically unassigned sequences and reveal protist lineages for which there is a major scarcity of taxonomic references, (ii) highlight unassigned protist diversity shared between contrasted marine environments and (iii) identify unassigned taxa which are ecologically relevant and recurrent, that should be prioritized for further characterisation. We integrated 12 years of data and 155 different sampling locations from 6 environmental metabarcoding datasets, combining 3 coastal time-series (ASTAN, BBMO, SOLA), 1 European coastal Sea sampling project (BioMarKs) and 2 oceanographic campaigns (Malaspina, MOOSE). As a study case, we focused our analyses on the parasite group of Syndiniales and, by clustering the gathered metabarcodes in a Sequence Similarity Network (SSN), we revealed novel ecological patterns of Syndiniales at a taxonomic resolution of 100% similarity between V4 regions of the 18S rDNA marker.

## Results

### Diversity and abundance of taxonomically unassigned protists: the uncharted territory of Syndiniales

Among the 343 165 metabarcodes we considered in our study (Table S1), those that were taxonomically unassigned at a given taxonomic rank (i.e., without any match with reference sequences under 80% of sequence similarity) according to the PR2 or SILVA reference databases were considered as unassigned at this taxonomic rank (Fig. S1A). Unassigned metabarcodes occured in every sampled region and at every taxonomic rank, from kingdom to species (Fig. 1A, Fig. S2). Both the relative abundance and number of unassigned metabarcodes increased from high to low taxonomic ranks contributing respectively to an average of 0.03% and 0.28% of the whole protist community at the kingdom rank and to 69.35% and 82.67% at the species rank (Fig. 1A, Fig. S3B). At kingdom level, 628 metabarcodes remained unassigned among which 87.70% originated from bathypelagic samples (2 150 - 4 000 m) of the Malaspina expedition (Fig. S4). The biggest increase in unassigned metabarcode proportion was observed from family to genus level for which 71.14% and 58.95% of metabarcodes were unassigned in relative number and relative abundance respectively (increase of 35% in unassigned metabarcodes). Overall, at the lowest taxonomic levels of our global dataset, i.e. genus and species, the proportion of unassigned metabarcodes was similar and represented more than half of the metabarcodes that could not be assigned to any referenced protist taxon (Fig. 1A). The study of unassigned sequences was thus conducted from the viewpoint of the genus taxonomic level.

**Fig. 1:**
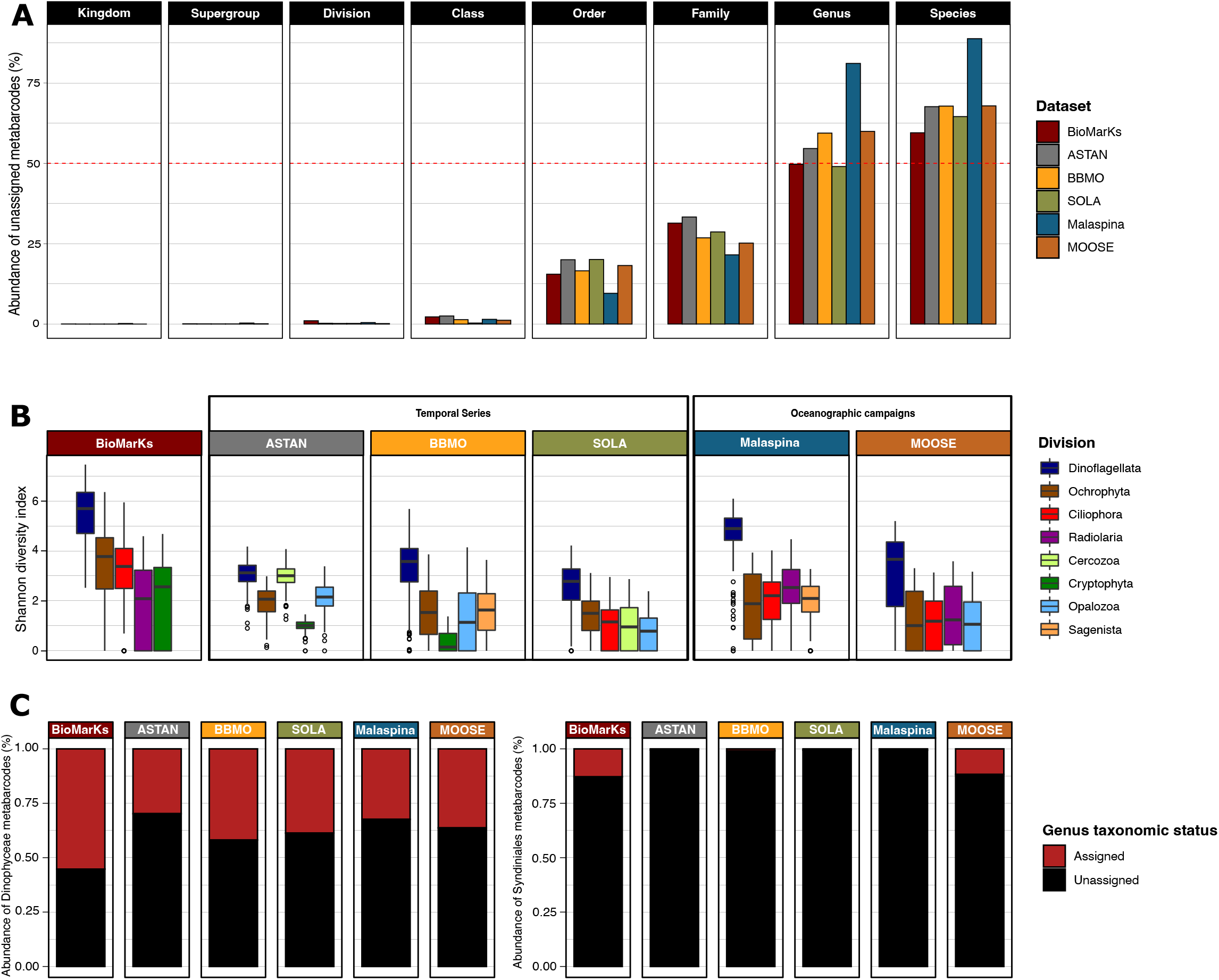
Relative abundance and diversity of unassigned metabarcodes. (A) Relative abundance of unassigned metabarcodes at each taxonomic level from kingdom to species. Colors represent the 6 studied datasets. The horizontal red dashed line marks 50% of the dataset in terms of relative abundance. (B) Shannon Weiner diversity index calculated at genus level within major protist divisions in each dataset. Only metabarcodes unassigned at genus level are selected. Colors indicate the protist divisions that represent >50% of unassigned metabarcodes at genus level in each dataset (Fig. S5). (C) Relative abundance of assigned and unassigned metabarcodes within the class Dinophyceae (left) and Syndiniales (right) found in each dataset. Colors indicate the taxonomic status (Assigned/Unassigned) of metabarcodes at genus level.

Across protist divisions, a higher diversity index was obtained for unassigned metabarcodes belonging to Dinoflagellata for all datasets (Fig. 1B). Overall, 54% of unassigned metabarcodes in relative number and 63% in relative abundance belonged to Dinoflagellata (Fig. S4A). Among other protist divisions lacking taxonomic assignment at the genus level were Ochrophyta (all datasets), Ciliophora (BioMarKs, SOLA, Malaspina, MOOSE), Radiolaria (BioMarKs, Malaspina, MOOSE), Cercozoa (ASTAN, SOLA), Cryptophyta (BioMarKs, ASTAN, BBMO), Opalozoa (ASTAN, BBMO, SOLA, MOOSE) and Sagenista (BioMarKs, BBMO, Malaspina) (Fig. S5A). A higher diversity index was obtained for unassigned sequences, compared to assigned sequences, for the divisions Opalozoa, Sagenista and Cercozoa (Fig. S5B). Thus, when studying only assigned genera of the latter protist divisions, their diversity could be largely underestimated.

Dinoflagellata metabarcodes represent 52% of our global dataset (179 615 metabarcodes). Among unassigned Dinoflagellata, Dinophyceae and Syndiniales were the two dominant classes and Syndiniales represented 66% and 48% of metabarcodes in terms of number and abundance respectively (Fig. S6A). Within these two classes, the proportion of unassigned metabarcodes at the genus level was 2-fold higher for Syndiniales, with 98% and 95% of metabarcodes unassigned in terms of relative number and abundance (Fig. 1C, Fig. S6B). Only 4 species of Syndiniales had a taxonomic assignment (0.01% of total metabarcodes and 0.53% of Syndiniales metabarcodes). Syndiniales metabarcodes unassigned at genus level represented 21% of our global dataset (72 789 metabarcodes). Given the contribution and overwhelming majority of unassigned Syndiniales in our dataset, we decided to focus the rest of our study on this lineage.

### Shared patterns of unassigned Syndiniales diversity between sunlit Mediterranean and Tropical waters

To investigate the spatio-temporal distribution of Syndiniales at genus level we built connected components (CCs), i.e clusters of metabarcodes with 100% sequence identity and a minimum of 80% coverage. We consider the CCs as a proxy for clustering metabarcodes of the same Syndiniales genera or at least as pragmatic units to deal with Syndiniales molecular diversity across multiple datasets. After clustering, our global dataset contained 4 317 Syndiniales CCs (30% of all CCs) out of which 4 245 CCs were unassigned at the genus level (98% of Syndiniales CCs) (Fig. S7A). These unassigned CCs belonged to 5 orders of Syndiniales, Dino-Group-I to III, Dino-Group-V and an “Unknown” order (rank not assigned). Out of the unassigned Syndiniales CCs, 58% (2 478 CCs) were exclusively shared within 2 sea regions, being mainly the Tropical/Subtropical Ocean and the Mediterranean Sea (which both include samples at depth > 1 000 m), regrouping 51% of the unassigned Syndiniales CCs (2 171 CCs) (Fig. 2, N=2). Unassigned CCs endemic to one region represented 23% of Syndiniales CCs (961 CCs) and were mostly found at the surface of the Tropical/Subtropical ocean (Fig. 2, N=1), while 12% of CCs (518 CCs) were shared between 3 regions (Fig. 2, N=3) and 7% CCs (288 CCs) were shared between more than 3 regions (Fig. 2, N>3). All studied sea regions shared 25 ubiquitous unassigned Syndiniales CCs, including 14 CCs belonging to the Dino-Group-II Syndiniales order (Fig. 2, N=6).

**Fig. 2:**
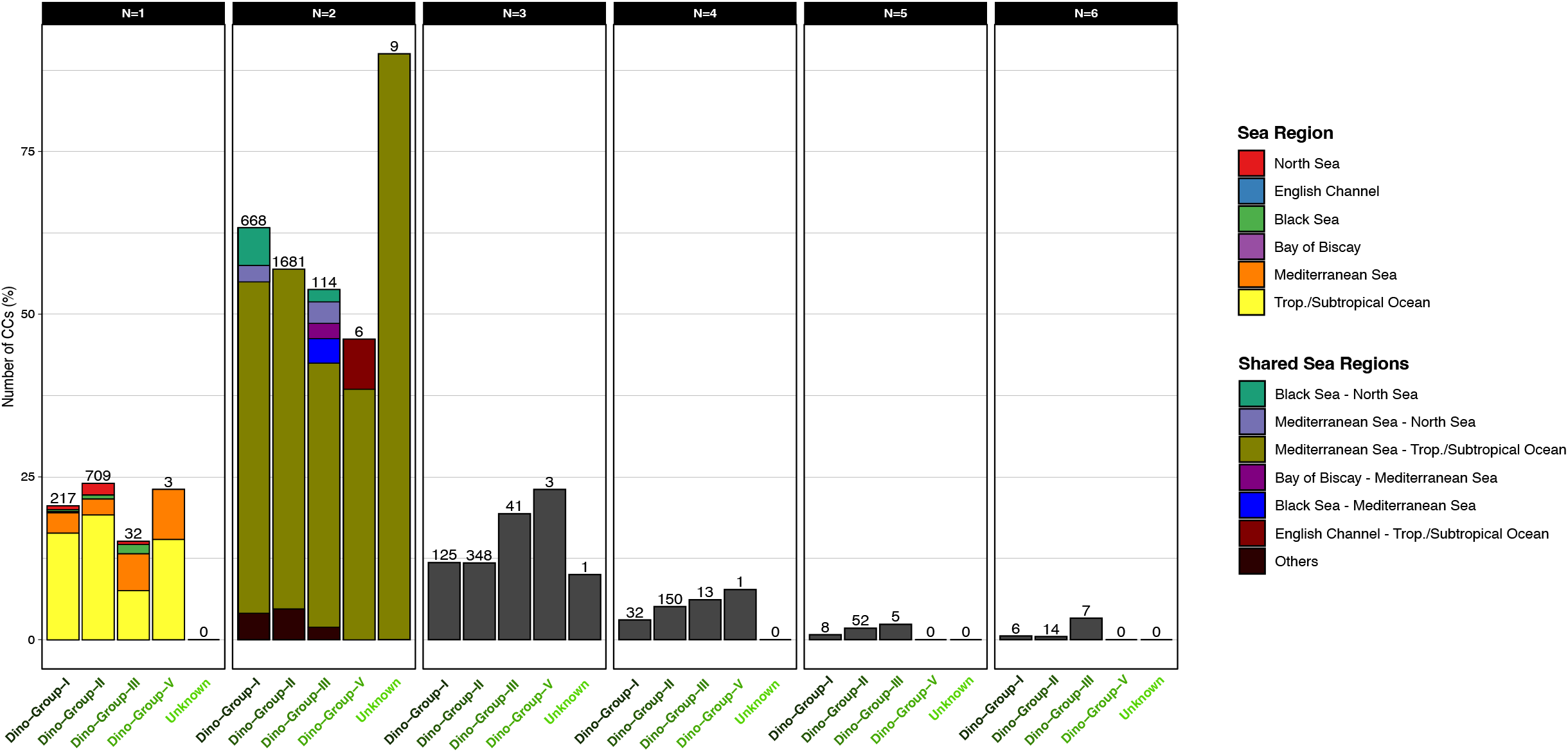
Distribution range of Connected Components (CCs) among Syndiniales orders unassigned at genus level. The relative number (%) of CCs (y axis) within each Syndiniales order (Dino-Group-I to V and Unknown, i.e. unassigned order) is represented according to their occurrence across the 6 sea regions defined in our metadata (Fig. S2). The raw number of CCs is indicated above each bar. The 4 245 CCs containing only unassigned sequences at genus level within each order of Syndiniales were selected. As Syndiniales order Dino-Group-IV contained only assigned sequences (at genus level) it was not included in the plot. Results are grouped on the x axis by the number of sea regions (defined by the PCA, Fig. S2) across which a CC is found (N). The colors indicate sea regions (for N=1) and pairs of sea regions (for N=2). The combination of sea regions is not illustrated for N>3. For N=2, “Others” include pairs of regions representing < 3.5% of pairs within each order. (N.b. The number of sampled stations is variable between sea regions with a higher number of stations sampled in Subtropical ocean (122 stations) and Mediterranean sea (35 stations). The other 4 sea regions are represented by samplings at a single station. Also Subtropical ocean and Mediterranean sea include samplings located between 200 m and 4000 m deep. North sea and Black sea samples are from surface, DCM and anoxic layers. The English Channel and Bay of Biscay include only surface samples (Table S1).)

Among Syndiniales orders, Dino-Group-II and Group-I were the most represented in our dataset (2 954 CCs, 70%; 1 056 CCs, 25% of unassigned Syndiniales CCs respectively (Fig. S7B)) and their distribution was mostly restricted to the Subtropical Ocean and Mediterranean Sea (Fig. 2). Dino-Group-III (212 CCs, 5% of unassigned Syndiniales CCs (Fig. S7B)) had the widest distribution, including diversity shared between many different pairs of regions with some patterns being unique to this order, i.e CCs that were exclusively common between the Bay of Biscay and the Mediterranean Sea and between the Black Sea and the Mediterranean Sea (Fig. 2, N=2). Dino-Group-V included 13 CCs (0.3% of unassigned Syndiniales CCs (Fig. S6)) and included CCs exclusively shared between the English Channel and the Tropical/Subtropical Ocean (Fig. 2, N=2). The Unknown Syndiniales order included 10 CCs (0.2% of unassigned Syndiniales CCs (Fig. S7B)) and was found in 3 sea regions: Mediterranean Sea, Tropical/Subtropical Ocean (main pattern for this order, 9 CCs) and North Sea (1 CC shared between the 3 mentioned sea regions) (Fig. 2).

Since for all Syndiniales orders 50% of unassigned CCs were found to be exclusively common to mediterranean and tropical regions we further explored how this pattern was distributed across the water column. Among the 2 171 CCs exclusively shared between mediterranean and tropical waters, Syndiniales communities were the most similar in the photic zone with 63% of CCs common between DCM (Deep Chlorophyll Maximum) layers and ∼30% common between surface and DCM reciprocally (34% CCs common between Tropical/Subtropical Ocean DCM and Mediterranean Sea surface; 32% CCs common between mediterranean DCM and Tropical/Subtropical Ocean, n.b. percentages are indicative of major trends and not proportion as combinations are not exclusive) (Fig. 3A). Notably, a pattern of shared Syndiniales CCs was also found between bathypelagic samples from the Mediterranean Sea and samples from the photic zone of the Tropical/Subtropical Ocean (29% CCs) (Fig. 3A).

**Fig. 3:**
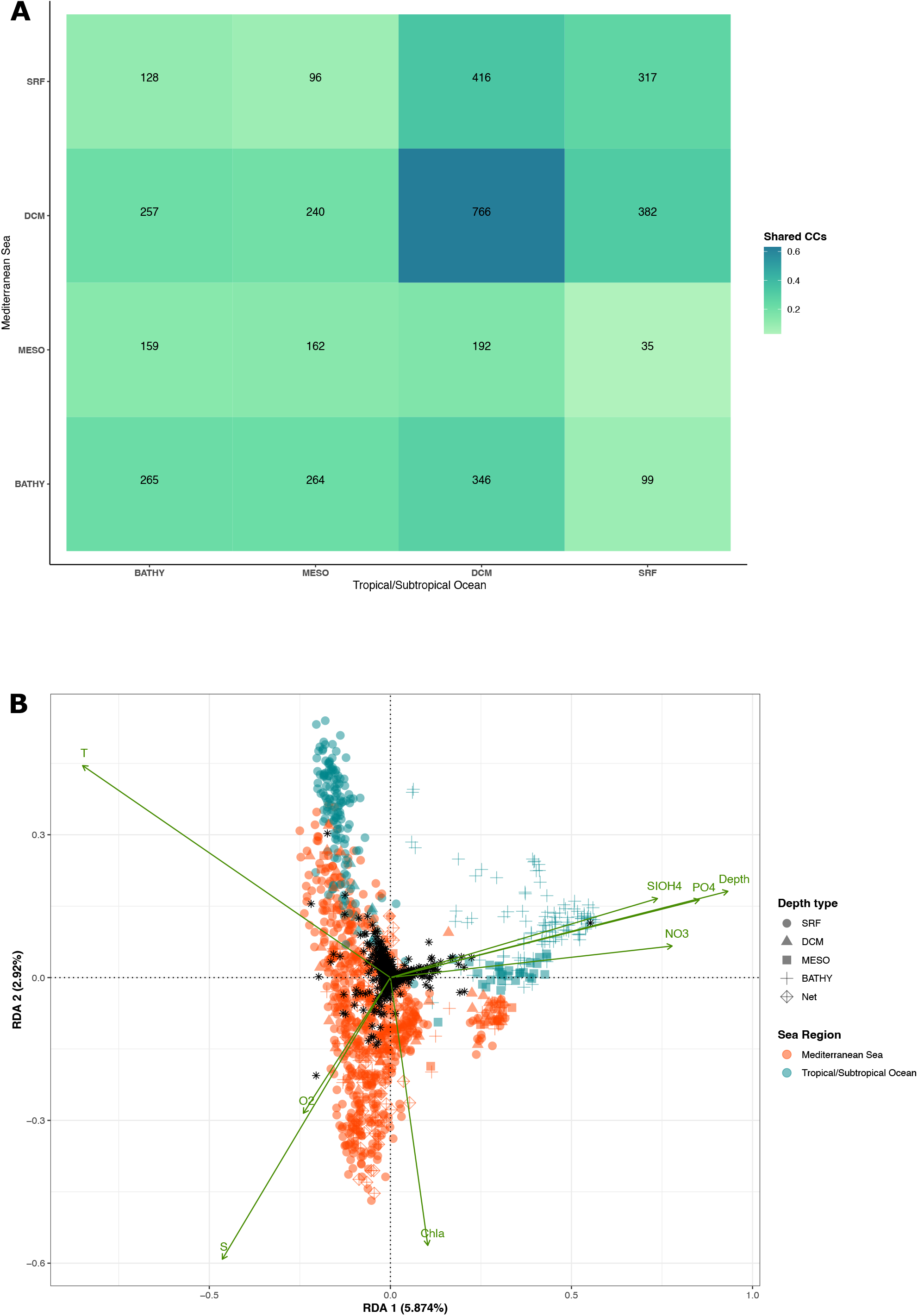
Similarity in Syndiniales genera communities between Mediterranean sea and Subtropical ocean. (A) Proportion of Syndiniales CCs unassigned at genus level and shared between the Mediterranean sea (y axis) and the Subtropical ocean (x axis) per depth layer (SRF for surface, DCM for Deep Chlorophyll Maximum, MESO for mesopelagic layer (>200-1 000m), BATHY for bathypelagic layer (>1 000-4 000m)). The percentages illustrate major trends and not proportions (i.e. sums of percentages exceed 100% as combinations of shared CCs are not exclusive and some CCs are present in multiple depth layers). The number of samples from each depth layer for the Mediterranean sea are: SRF=571, DCM=88, MESO=97 and BATHY=46 and for the Subtropical ocean: SRF=136, DCM=13, MESO=30 and BATHY=110. The number of CCs found in each depth layer is: SRF=1 620, DCM=1 221, MESO=449 and BATHY=518 for the Mediterranean sea and SRF=1 726, DCM=943, MESO=611 and BATHY=1 281 for the Subtropical ocean. (B) Redundancy Analysis (RDA) for Mediterranean Sea and Subtropical Ocean data. The variation of abundance in unassigned Syndiniales CCs (black stars) is correlated to the variation of physicochemical parameters (green arrows). The most pertinent environmental parameters allowing to differentiate the studied marine regions were selected (cf. Materials and Methods: Spatiotemporal patterns of metabarcodes and CCs). The samples are represented by different colors for Mediterranean Sea (orange) and Subtropical Ocean (blue). The shapes indicate the depth layer: dot; SRF, triangle; DCM, square; MESO, cross; BATHY and square/cross; NET (vertical profile samples (0-500m)). The dimensions of the input abundance matrix are: 4 037 CCs and 1 055 samples (768 samples for the Mediterranean Sea and 287 for the Subtropical Ocean). The global RDA (cf. Materials and Methods: Spatiotemporal patterns of metabarcodes and CCs) was statistically significant at 0.005% and the first 2 axes of the RDA with the selected explanatory variables (shown below) were significant at 0.01%.

In order to test if these shared diversity patterns can be explained by similar physicochemical conditions, we explored the abundance variation of unassigned Syndiniales CCs in the mediterranean and tropical waters in an RDA using the physicochemical parameters as explanatory variables. The variation of physicochemical parameters explained ∼10% (11.2% in first 6 RDA dimensions) of the abundance variation of unassigned Syndiniales CCs (Fig. 3B). Based on this result, two communities of Syndiniales could be distinguished according to the first two dimensions of the RDA: a deep water (> 200 m) community associated with colder and more eutrophic conditions (Fig. 3B, left) and a photic (surface and DCM) community associated with warmer and more oligotrophic conditions (Fig. 3B, right). In the RDA space associated with the photic zone, mediterranean and tropical samples partly overlap and correspond to warmer and less salty waters, hence providing an environmental basis for the observed Syndiniales pattern in these two marine environments (Fig. 3A).

To further investigate this hypothesis, we compared the community composition of protist divisions known to be major hosts for Syndiniales, between the marine regions of our global dataset. For Dinophyceae, Radiolaria and Ciliophora, the jaccard dissimilarity index was the lowest between Mediterranean sea and Tropical/Subtropical Ocean compared to community comparisons between the other sea regions (Table S2). This was also the case for Syndiniales, further supporting the results illustrated above (Fig. 3A). Neither of the remaining protist divisions found as having an important contribution to our dataset (Ochrophyta, Cercozoa, Cryptophyta, Opalozoa) showed the same tendency apart from Sagenista (Fig. S4, Table S2).

### Rhythmic ecological indicators among unassigned Syndiniales community

Temporal aspects of the Syndiniales community were studied across the three time-series (ASTAN, BBMO, SOLA) in our dataset. Unassigned Syndiniales clusters did not indicate any clear seasonal preference based on monthly abundance for any of the time-series (Fig. S8). The correlation of CCs to the overall Syndiniales community dynamics and their rhythmicity was computed with two methods. The Escouffier’s equivalent vectors selected the CCs that are the best indicators of community abundance variation according to a PCA and the Lomb-Scargle periodogram algorithm detected if CCs follow rhythmic patterns of occurrence across time. In the studied time-series, 75% of the Syndiniales community response to environmental variation was described by 45 CCs at ASTAN, 36 CCs at BBMO and 17 CCs at SOLA (Table S3). These community indicator CCs were all unassigned at the genus level. Rhythmic occurrence among Syndiniales CCs was found to be more prevalent in the Western Channel with 208 rhythmic Syndiniales CCs found at ASTAN, 118 CCs found at BBMO and 15 CCs found at SOLA (Table S4). Some of the unassigned Syndiniales CCs were found to be both community indicators and rhythmic throughout the time-series: 27 CCs at ASTAN, 7 CCs at BBMO and 5 CCs at SOLA (Table S5). The average recurrence period of these clusters was ∼1.5 years at ASTAN and BBMO ∼1 year at SOLA (Table S6). We identified two rhythmic indicator CCs shared between the time-series of the English Channel (i.e. ASTAN) and Mediterranean Sea (Fig. 4): CC_unknown_154, shared with BBMO, and CC_unknown_183, shared with SOLA (recurrence periods are indicated in Table S6). One indicator CC, CC_unknown_126, was found to be shared between all the studied time series (Fig. 4) with quicker recurrence periods in the Mediterranean (Table S6). All other rhythmic indicator CCs were specific to each time-series. CC_unknown_126 was the CC with the highest monthly relative abundance at BBMO and SOLA, while having the 4th highest monthly relative abundance at ASTAN. The seasonal prevalence for the majority of rhythmic indicator CCs was up to 3 seasons (Fig. 4, Table S7). Rhythmic indicators with a 4 season prevalence occurred and were more numerous at the Western Channel. The shared indicator CC_unknown_126 maintained a high seasonal prevalence occuring at 3 seasons in the Mediterranean Sea (i.e., BBMO and SOLA) and 4 seasons in the English Channel (i.e., ASTAN) (Fig. 4).

**Fig. 4:**
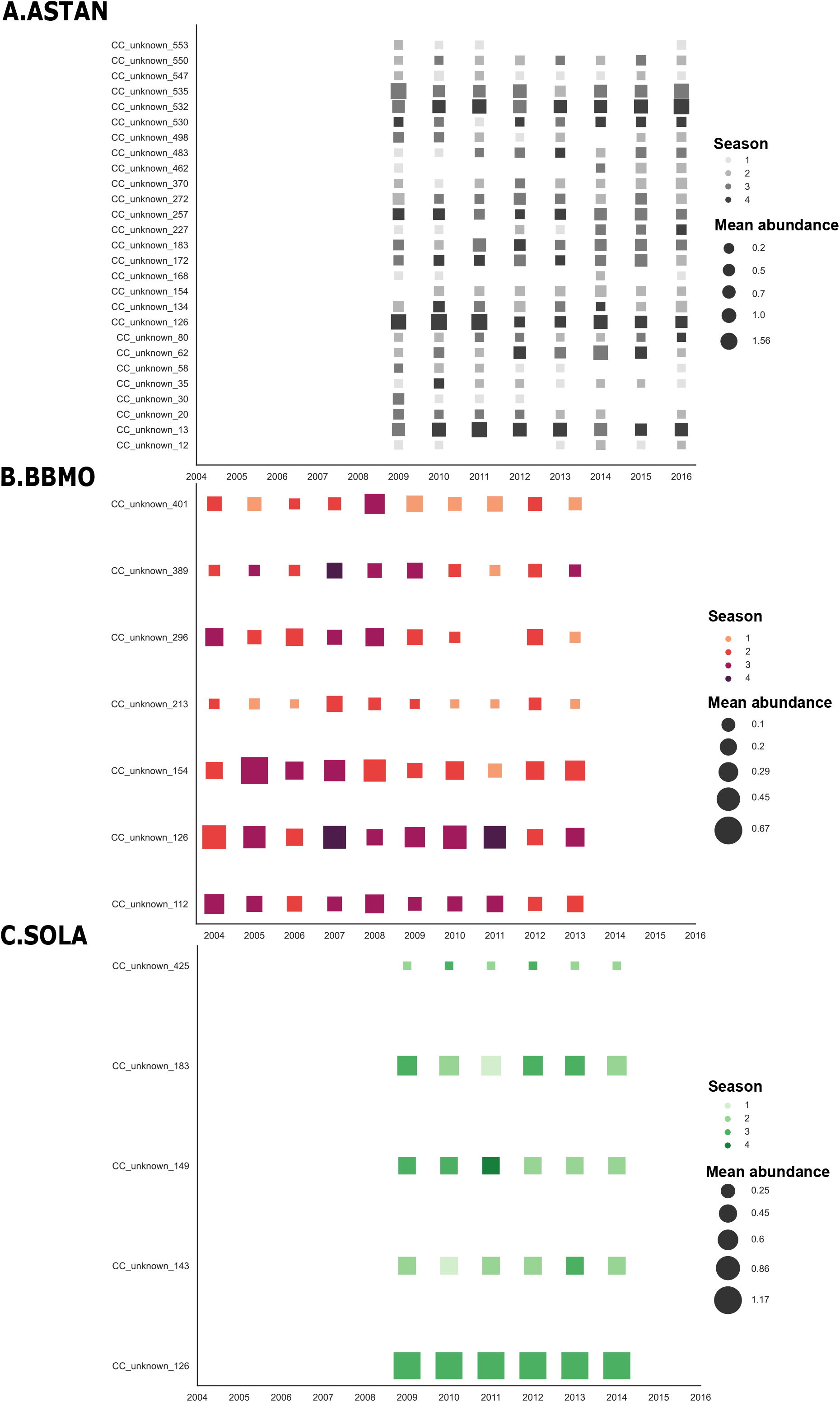
Annual seasonal prevalence and abundance of rhythmic indicator Syndiniales CCs. The occurrence of CCs selected by the Escouffier’s equivalent vectors and Lomb-Scargle Periodogram methods was studied across each time-series: (A) ASTAN; (B) BBMO; (C) SOLA. Relative abundance was computed per year as an average value of each month and is represented by square size. Colors indicate the seasonal prevalence of the CC throughout each year and the color gradient indicates the prevalence extent (i.e. 1 season prevalence indicated by the lightest color and 4 seasons indicated by the darkest color of the gradient). A CC is considered prevalent if it is present at least once during each season. Taxonomically unassigned CCs at genus level are indicated by “unknown” in the CC ids (y axis).

## Discussion

### What are we missing from eukaryotic diversity with metabarcoding ?

In environmental genomics investigations, the 18S rDNA universal marker sequence constitutes the gold standard for the exploration of eukaryotic diversity in environmental communities, shedding light on uncultivable and rare taxa [16, 30]. Yet, by integrating different metabarcoding datasets, we report that in the marine realm half of protist sequences cannot be taxonomically assigned at the genus level (57% of sequences in our dataset) and these unassigned protist taxa represent 36% to 82% of the protist community in terms of abundance across 6 diverse marine environments. Few metabarcoding studies have quantified unassigned protist diversity. In Tara Oceans, unassigned protist diversity revealed with the V9 region of the 18S rDNA marker at the supergroup level was found to be <3% of total reads [14] when referring to unassigned sequences as marker sequences with <80% identity with reference sequences. Here, with the V4 region of the 18S rDNA marker we find that unassigned protist sequences represent in abundance <1% at the supergroup level. At the genus level unassigned sequences are not rare among the protist community as they represent in abundance >45% of metabarcodes in each studied dataset and up to 80% of metabarcodes for the Malaspina expedition dataset.

This confirms the current biased view of eukaryotic diversity, mostly focusing on multicellular and cultivable taxa, neglecting >70% of eukaryote diversity, including key lineages for the evolution of life and to understand ecosystems functioning [30-32]. This missing picture can be addressed, for metabarcoding studies, in the context of sample acquisition but also data acquisition in reference databases. Some oceanic regions are more sampled than others, i.e. coastal locations compared to deep / open-sea environments [33]. Moreover, the maintenance and update of reference databases is a laborious but critical process whose pace is difficult to synchronize with the generation of an ever-increasing amount of environmental sequences [34]. Metabarcoding assessments of the diversity also depends on the choice of ribosomal marker genes. In our study, the largest proportion of unassigned protist diversity was found at low taxonomic levels, a trend that has also been observed for prokaryotes [6]. Universal ribosomal markers such as 16S rDNA and 18S rDNA can have a distinct taxonomic resolution depending on the lineage considered and within each lineage [30, 35], for instance in order to describe diatom diversity a threshold >95% similarity of the V9 regions of the 18S rDNA gene with reference sequences delimits some genera (e.g. Undatella) while a threshold of <90% is sufficient for assigning some other genera (e.g. Synedropsis) [36].

### Perspectives on Syndiniales biogeography

The challenge of barcoding marker taxonomic resolution is particularly relevant for rapidly evolving lineages, like predicted by evolutionary theory for parasites [37]. Studies on life history traits of multicellular parasites have demonstrated their quick adaptive plasticity, being involved in an evolutionary arms race with their host [37, 38]. Parasites are the most abundant component in many eukaryotic communities investigated through metabarcoding approaches, whether using high throughput sequencing technologies such as Illumina in tropical soils [40], subtropical marine ecosystems [39] and polar regions [20, 21], or low throughput cloning-sequencing methods in a lacustrine ecosystem [41]. In our study, parasitic Dinoflagellates (Syndiniales) represented 22% of metabarcodes and only 0.4% (1 537 metabarcodes) could be assigned to a referenced genus, being the major contributor to the unassigned marine protist microbiome. To capture efficiently Syndiniales diversity, an alternative would be to design specific primers as it has been done to target Perkinsea and Microsporidia [43], or a combination of distinct genetic markers should be favored (e.g. 18S and ITS or COI) [43]. The cytochrome *c* oxidase 1 (COI) barcode has successfully identified different cultured dinoflagellate species [44]. Studies using the V9 and V4 regions of 18S rDNA marker [45] to study the diversity of Dinoflagellata, retrieved different diversity patterns for each marker, stressing the difficulty to describe diverse, dominant lineages of protist communities.

When studying the distribution patterns of Syndiniales, we found CCs of the 100%-similar sequences shared between disconnected oceanic regions included along a latitudinal gradient from the North Sea to the South Subtropical Atlantic, Indian, and Pacific Oceans. Clarke *et al*., 2019 have reported an OTU from Syndiniales Group I with identical V4 regions of the 18S rDNA marker to have been retrieved from surface samples in a Southern Ocean transect near sea-ice edge and seven different Northern Hemisphere coastal locations including tropical/subtropical zones. The putative inferred V9 region of 18S rDNA marker of this abundant Syndiniales is present in every station of the Tara Oceans voyage, including mediterranean samples [20]. This suggests that closely related parasites can infect a wide range of hosts [20], which could also be the case for the shared Syndiniales CCs in our study. Our results indicated 50% (2 171 CCs) of the Syndiniales community in common between two tropical/subtropical waters and the mediterranean basin in the euphotic zone. In this case, a convergent selection of host-parasite systems in distant but physicochemically similar oligotrophic environments could also be hypothesized. Statistical analyses reported physicochemical similarities for surface waters of these marine environments and potential host communities showed greatest similarity in composition between the Tropical/Subtropical Ocean and the Mediterranean Sea. In that context, the shared Syndiniales CCs between bathypelagic tropical/subtropical and photic mediterranean layers could be linked to different life stages of hosts across the water column [46]. Complementary studies need to be done comparing open sea and coastal regions, using alternative markers to 18S rDNA to validate these observations. Further exploring host-parasite comparative biogeography patterns through co-occurrence networks could help elucidate the distribution extent of parasite associations at a global scale and allow to define more precisely host-ranges among parasites at low taxonomic resolution [47-49].

### Perspectives on Syndiniales temporal dynamics

By studying temporal patterns of Syndiniales across 3 time-series we highlighted a small number of CCs that are recurrent over time, persistent through seasons and some indicators of parasite community variation. The recurrence of these taxa could be associated with rhythmic host patterns like annual blooms, as parasites can respond quickly to elevated host density [22, 26]. Taxa persistent throughout seasons could further indicate a generalist and opportunistic parasite behavior, infecting the hosts that are present during each season, while surviving in spore form during low host densities [23, 50]. Flexible host-parasite associations have already been described in coastal estuaries using co-occurrence networks [22]. Alternatively, parasites cannot persist below a critical host threshold [28], which questions the trophic mode of the detected persistent Syndiniales. Up to date only parasitic and parasitoid Syndiniales have been described [47]. Nevertheless, parasitism is a mode of symbiosis along a parasite-mutualist continuum and transitions from one mode to the other should not be excluded [51].

### Syndiniales as potential indicators of ecosystem change ?

Our analysis also highlighted Syndiniales CCs that were both recurrent over time and good indicators of parasite community abundance variation. These Syndinales CC hold the potential for monitoring changes in environmental microbial communities, reflecting shifts not only among the Syndiniales communities but also mirroring their associated host community. The absence of these Syndinales CC could, for instance, indicate a shift in microbial community composition during or after an environmental perturbation. For instance, in marine environments, multicellular parasites (e.g. trematodes) have been employed as bioindicators of host physiology in response to accumulating pollution for environmental monitoring [52]. The diversity of frog and fish endoparasites was shown to reflect their surrounding ecological conditions. Selecting endoparasite taxa that are sensitive to environmental perturbation is crucial for a potential bioindicator. In that respect, our analysis throughout a 6-10 years of abundance information and metadata suggest that dinoflagellate parasites could be used for marine habitat monitoring as it has been done with diatoms, ciliates and foraminifera [30]. Behind the blackbox of Syndiniales taxonomy could be hidden a promising global ecosystem change indicator; thanks to their worldwide distribution [14], abundance [22], quick response time to host community shifts (Anderson & Harvey 2020) and intimate implication in marine food webs [26].

### SSNs as integrative tools to prioritize unassigned protist taxa

In this integrative study we have used a sequence similarity network to explore the ecology of the main components of the unassigned protist microbiome by combining 6 metabarcoding datasets. SSNs are relevant and efficient analytical tools for addressing the unassigned microbiome challenge as they allow studying simultaneously large datasets, in order to categorize and prioritize unassigned sequences. They have been recently employed among prokaryotes for surveying the coding part of genomes and metagenomes [6] and taxonomy across extreme aquatic environments [12]. By exploring the biogeography of these sequences we can reveal core taxa shared across ecosystems [12, 53]. Here we have explored both biogeographical and temporal patterns of protists at the species level without requiring a reference taxonomic match. Our FAIR (Findable, Accessible, Interoperable and Reusable) computational workflow that allows to integrate data from heterogeneous ecosystem sampling protocols, such as coastal time-series and open sea campaigns and can be applied to any targeted protist group of any metabarcoding dataset, of the same marker gene, for example originating from the metaPR2 database [54] and Ocean Barcode Atlas [55]. The taxa identified by the network could then be specifically targeted for *in situ* hybridisation [43] and isolation for single-cell omics [31]. Other approaches to reduce the unassigned taxonomic load encompass long-read sequencing [16], sequencing multiple metabarcoding markers [30] and combining metabarcoding and microscopy [34]. The unassigned microbiome holds an unexplored potential of novel taxa and functions that will surely challenge the current view of microbial ecology in the ocean and beyond [5, 31, 56].

## Materials and Methods

### Gathering and homogenisation of metabarcoding datasets

Metabarcoding datasets of 18S rDNA marker sequences containing the variable region V4 and originating from 6 distinct sampling projects were gathered. The datasets include three temporal series of bimensual samplings at a single station: ASTAN in Roscoff, English Channel, France (8 years of data), BBMO in Blanes Bay, Mediterranean Sea, Spain (10 years of data) and SOLA in Banyuls-sur-Mer, Mediterranean Sea, France (9 years of data) (Fig. S1D); two oceanographic campaigns of punctual samplings across 148 locations: Malaspina Expedition (122 stations, circumglobal Tropical/Subtropical Ocean) and MOOSE (26 stations, Mediterranean Sea) (Fig. S1B,C); and one European project of punctual samplings at 6 marine coastal stations: BioMarKs project (samples from: Oslo, Norway; Roscoff, France; Varna, Bulgaria; Gijon, Spain; Barcelona, Spain; Naples, Italy). Sequencing was done with Illumina MiSeq technology, except for the BioMarKs project sequenced by 454 pyrosequencing. Each metabarcoding dataset contained the abundance tables of reads clean-processed and inferred into ASVs (OTUs for BioMarKs) and their taxonomic affiliation (details in Table S1). The initial global dataset contained 539 546 metabarcodes. For homogenisation purposes, the same two filtering conditions were applied independently to each of the 6 datasets (Fig. S1A, Step 1): removal of sequences corresponding to metazoans, terrestrial plants (Streptophyta) and macroalgae (Florideophyceae, Bangiophyceae, Phaeophyceae, and Ulvophyceae); removal of sequences having less than 80% identity with reference databases. The latter threshold was chosen according to the original preprocessing of the datasets: the MOOSE dataset had beforehand implemented a minimum identity threshold of 80% and Malaspina and BBMO of 95%. A 95% filter was considered too stringent, as too many unknown sequences of interest might be removed, a 80% threshold was applied to the global dataset for homogenisation. The global abundance table resulting from the homogenisation workflow involved at this stage 343 165 metabarcodes, and each sample was normalized by total read number and scaled from 0 to 1.

To account for variations in the taxonomic assignment procedure (assignment tools, database versions) across datasets, a new taxonomic assignment (Fig. S1A, Step 2) was performed on the global set of metabarcodes with the PR2 database (version 4.12.0, released on 08.08.2019, https://pr2-database.org ; blast parameters: -evalue 0.01 -max_target_seqs 15, [57]). Only the best hit (best e-value) of each alignment was kept. These new assignments were filtered again for multicellular taxa and only sequences with a length greater to 200 bp were kept (Fig. S1A, Step 3). The PR2 database includes 8 taxonomic ranks: kingdom, supergroup, division, class, order, family, genus, and species. To avoid prokaryotic contamination at the kingdom level, an assignment was performed using the SILVA database (https://www.arb-silva.de/, version 138) implemented in the DADA2 algorithm [58]. 3 874 prokaryotic metabarcodes were removed out of the 4 519 unassigned sequences at the kingdom level. The taxonomic ranks that were left unassigned were marked as “Unknown” and the taxonomy of the sequence was considered unassigned at this given rank. Unassigned ranks located between attributed ranks were regarded as gaps in the taxonomic hierarchy and not as unassigned ranks. The diversity and abundance of unassigned sequences were explored on Rstudio (R version 4.1.1, [59]), using the packages: ‘data.table’, ‘vegan’, ‘ggplot2’, ‘ggsci’ and ‘gridExtra’.

### Homogenisation and analysis of environmental data

Our global dataset included 1 531 samples (ASTAN: 374, BBMO: 327, SOLA: 154, Malaspina: 289, MOOSE: 272) (Table S1). The metadata and environmental information associated with the studied samples were retrieved from the initial studies [60-66] and supplemented with public oceanographic databases (cf. additional information in the next paragraph). The information contained 14 metadata variables: name of the campaign, sampled region, station (for oceanographic campaigns), sequencing technology, sampling date, year, month, season, depth (m), depth type (surface (depth **:s** 5 m), deep maximum chlorophyll (DCM), mesopelagic zone (depth **2** 200 m), bathypelagic zone (depth **2** 1 000 m)), sampled size fraction, latitude, longitude). The 3 temporal series datasets (ASTAN, SOLA, BBMO) were sampled only at surface, BioMarKs dataset was sampled at surface and DCM, while the 2 oceanographic campaigns (MOOSE and Malaspina) were sampled at surface, DCM, mesopelagic and bathypelagic zones (up to 2 000 m depth for MOOSE and 4 000 m for Malaspina). The sampled size fractions are: 0-0.2 µm, 0.2-3 µm, 0.2-0.8 µm, 0.8-3 µm, 0.8-20 µm, 3-20 µm, 20-2 000 µm. The information contained as well 10 environmental variables: temperature (°C), salinity (PSU), pH, concentrations of oxygen (ml/L), nitrate (µmol/L), nitrite (µmol/L), ammonium (µmol/L), phosphate (µmol/L), silicate (µmol/L) and chlorophyll-*a* (µg/L). For ASTAN and BioMarKs datasets, when *in situ* environmental variables were missing, metadata were retrieved from public oceanographic databases (SOMLIT database (https://www.somlit.fr) ; World Ocean Database (http://www.ncei.noaa.gov/access/world-ocean-database-select/dbsearch.html) www.ncei.noaa.gov/access/world-ocean-database-select/dbsearch.html), SeaDataNet (https://cdi.seadatanet.org/search)). No additional information could be retrieved for 2 locations (Varna and Gijon). The environmental data and metadata were explored on Rstudio (R version 4.1.1), using the packages: ‘maps’, ‘tidyverse’, ‘sp’, ‘reshape2’, ‘tidyr’, ‘ade4’, ‘factoextra’ (Principal Component Analysis), ‘ggplot2’, ‘ggsci’ and ‘gridExtra’.

### Sequence Similarity Network as a framework for heterogeneous datasets comparison

The 343,165 metabarcodes were aligned against each other with the following options: e-value < 1e-4 ; >80% coverage for both subject and query (except for the alignments involving SOLA sequences (maximum sequence length = 230 bp compared to a mean of 430 bp for other datasets) in which case the coverage threshold was applied only to the SOLA sequence in order to avoid a misrepresentation of SOLA sequences in our analysis). Self-hits and reciprocal hits (same query-subject pair) were discarded.. The filtered blast output (2 942 982 alignments) was used to cluster sequences by similarity in a Sequence Similarity Network (SSN), with ‘igraph’ R package (version 1.2.6, https://igraph.org/r/, [67]). The sequences (i.e., the network nodes) were labeled according to metadata and taxonomic affiliation. The sequences were clustered into Connected Components (CCs) by setting an identity threshold of 100% sequence similarity, and CCs involving less than 6 sequences were removed (this number of 6 was chosen in order to enable the representativity of all 6 datasets in small CCs. The taxonomic homogeneity of CCs in the network was evaluated for known sequences at the genus level, and if only a single genus assignment was found this name was extrapolated to the other nodes of the CC even if these ones were of unknown genera. Thus, CCs were considered here as a proxy for studying taxonomic diversity at the genus level. The final network was composed of 12 619 CCs.

### Spatio-temporal patterns of metabarcodes and CCs

CCs including only Syndiniales sequences unassigned at genus level were extracted from the network (4 245 CCs; 33.6% of network and 47.6% of unassigned network CCs at genus level, Fig. S7A, Fig. S8). The distribution of clusters across marine environments and time was explored with R functions that were coded to extract the sequence attributes related to sampling data in each CC (location, dataset, depth, season month). A Redundancy Analysis (RDA) was performed on the abundance matrix of Syndiniales CCs using the metadata for Tropical/Subtropical Ocean and Mediterranean sea samples as explanatory variables. ANOVA tests were run to assess the robustness of the global RDA (all environmental variables included) and of the first two dimensions of the RDA with selected environmental variables. Both the RDA and ANOVA were run via the *vegan* package. Potential Syndiniales host communities were compared with the Jaccard dissimilarity index of the based on the Bray-Curtis compositional dissimilarity of abundances [68]. Jaccard index was computed with the *vegdist* function of the ‘vegan’ package, according to the formula: 2B/(1+B), where B is Bray-Curtis dissimilarity. The temporal patterns of Syndiniales among each Time Series (ASTAN, BBMO, SOLA) were explored for both assigned and unassigned genera clusters (4 317 CCs ; 34.2% of network, Fig S7B). Diversity indexes (species richness (S), Shannon’s diversity (H) and reverse Pielou index (J), using the *vegan* package) and statistical metrics (mean abundance per month) were computed. The Escoufier’s equivalent vector method was applied on CCs present at least 5 times across each time series. This method was run with the package *pastecs* and sorted clusters according to their correlation to a principal component analysis (PCA) [69]. The cumulated correlation level chosen was 75% in order to avoid retrieving clusters with negligible correlation (100% would result in retrieving the whole dataset). The rhythmicity of CCs across time was computed by the Lomb-Scargle Periodogram (LSP) [70] via the *lomb* package. Each CC was associated with a PNmax value, a p-value and a rhythmicity period (in days). The LSP method was applied according to Lambert et al., 2019 and is particularly well suited for our time-series data, as it allows us to detect the periodic patterns in unevenly sampled data. The PNmax is the decision variable corresponding to the peak normalized power, and CCs were considered rythmic for a PNmax > 10 (i.e. p-value < 0.01). Graphical representations were plotted on Rstudio (R version 4.1.1) and Python (v3.8, package ‘seaborn’).

## Acknowledgements

This work was supported by the *Institut des Sciences du Calcul et des Données* (ISCD) of Sorbonne University via funding from the project FORMAL (From ObseRving to Modeling oceAn Life, https://iscd.sorbonne-universite.fr/research/sponsored-junior-teams/formal-2/). The authors thank the researchers that provided the datasets and metadata for this work: N Simon and M Caracciolo (Ecology of Marine Plankton (ECOMAP), Station Biologique de Roscoff, France) for the ASTAN time series, M Mendez-Sandin (Systematic Biology, Dept. of Organismal Biology Uppsala University, Sweden) for the MOOSE campaign, R Logares (Institut de Cie’ ncies del Mar (ICM), Barcelona, Spain) for the Malaspina campaign, Blanes Bay Observatory (BBMO) time series and the BioMarKs project, P Galand (Laboratory of Microbial Oceanography, Banyuls-sur-Mer, France) for the SOLA time series. All analyses of this work were performed remotely on the AbiMs cluster of the marine station of Roscoff (http://abims.sb-roscoff.fr). L Bittner acknowledges the Institut Universitaire de France for her 5-year nomination as Junior Member (2020-2025).□

## Data accessibility

Scripts, data and Rmarkdown files necessary to run all the analyses included in this work are publicly available on the github page https://github.com/IrisRizos/Unassigned_Protists_SSN.

## Graphical Abstract

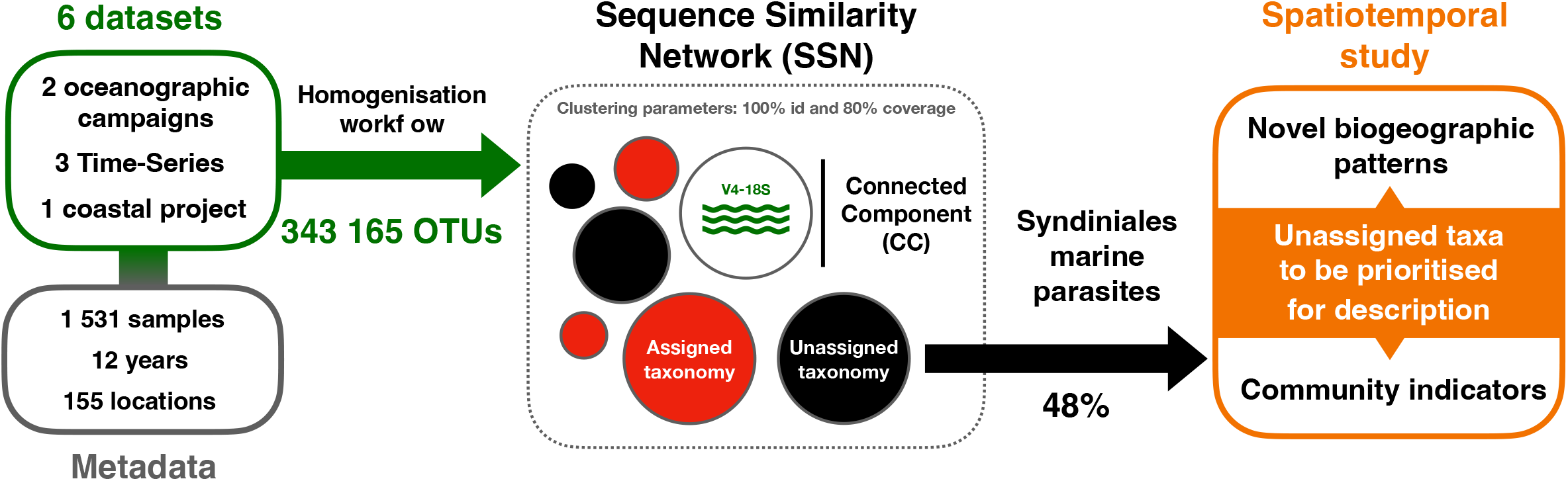

