## Supplementary Materials for "Beyond the limits of the unassigned protist microbiome: inferring large-scale spatio-temporal patterns of marine parasites"

#### Supplementary Figures

**Fig. S1:** The 6 metabarcoding datasets gathered for this study. (A) Summary of homogenisation workflow and analysis for integrating the metabarcodes from the 6 metabarcoding datasets. Unassigned sequences kept after the homogenisation workflow were first studied through α-diversity measures within each dataset and then combined in the Sequence Similarity Network for spatiotemporal exploration (B, C) Maps indicating the sampling sites of each dataset. The number of different sampled locations for each dataset is: (B) ASTAN 1, BBMO 1, MOOSE 26, SOLA 1 ; (C) BioMarKs 6, Malaspina 122. The total number of different sampled locations in the global dataset is 155. (D) Timeline of the sampling periods covered by each metabarcoding datasets. The cumulative years of data in the global dataset covered by frequent sampling of the time-series correspond to 12 years of data. BioMarKs, Malaspina and MOOSE datasets include punctual samplings at the indicated years.

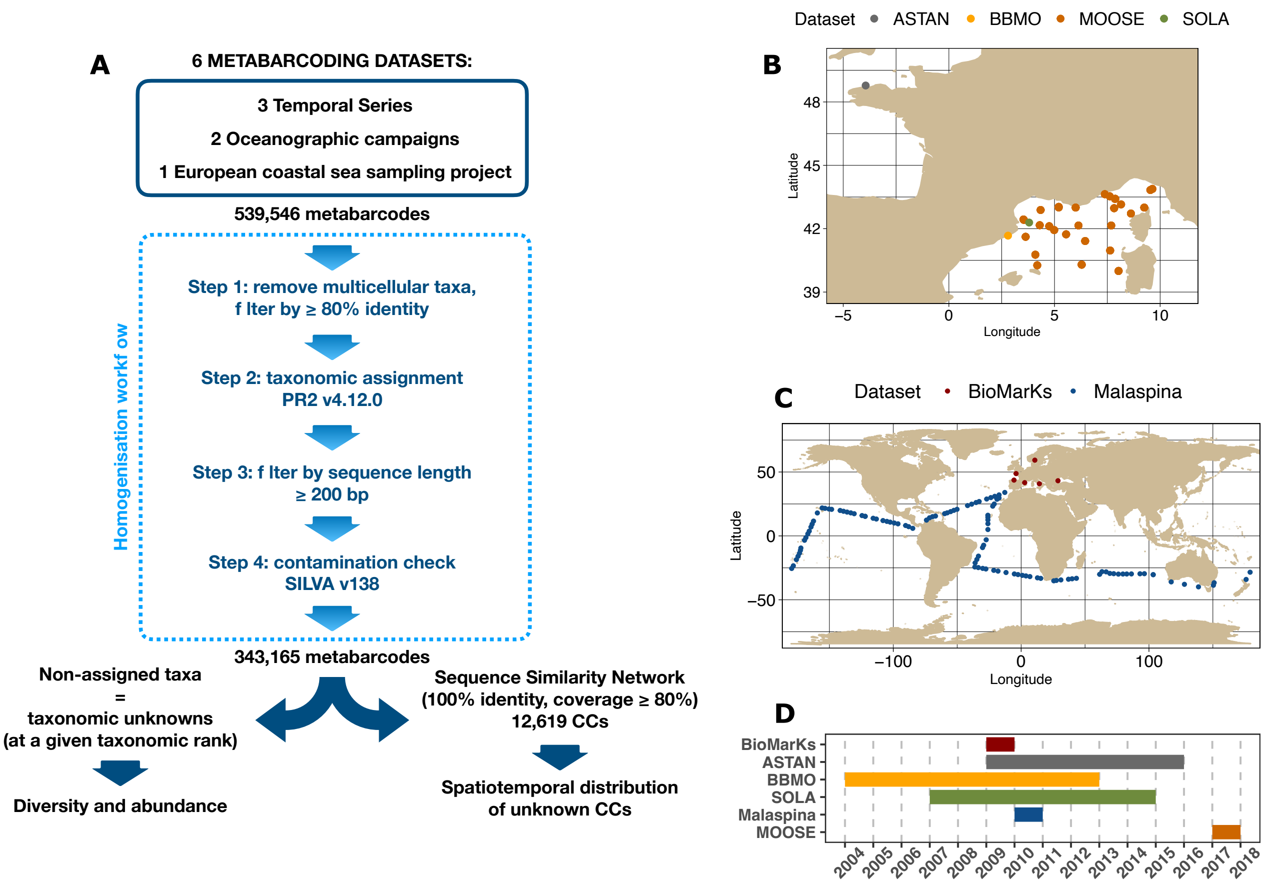

**Fig. S2:** Principal Component Analysis based on the metadata of the 6 studied metabarcoding datasets. PCA biplot with samples and physicochemical parameters (left): Chla (Chlorophyll α), $O_2$ (dissolved oxygen), Latitude, Longitude (spatial coordinates of the sampling spot), $NO_2$ (nitrate), $NO_3$ (nitrite), $NH_4$ (ammonium), $PO_4$ (phosphate), $SIOH_4$ (silicate), Depth, T (temperature), pH, S (salinity). The first 2 axes of the PCA explain 44.6% of metadata variation. PCA individuals (right) with samples colored by marine region. According to the first 2 axes of the PCA samples are grouped based on geographical location and depth, delimiting 6 marine regions.

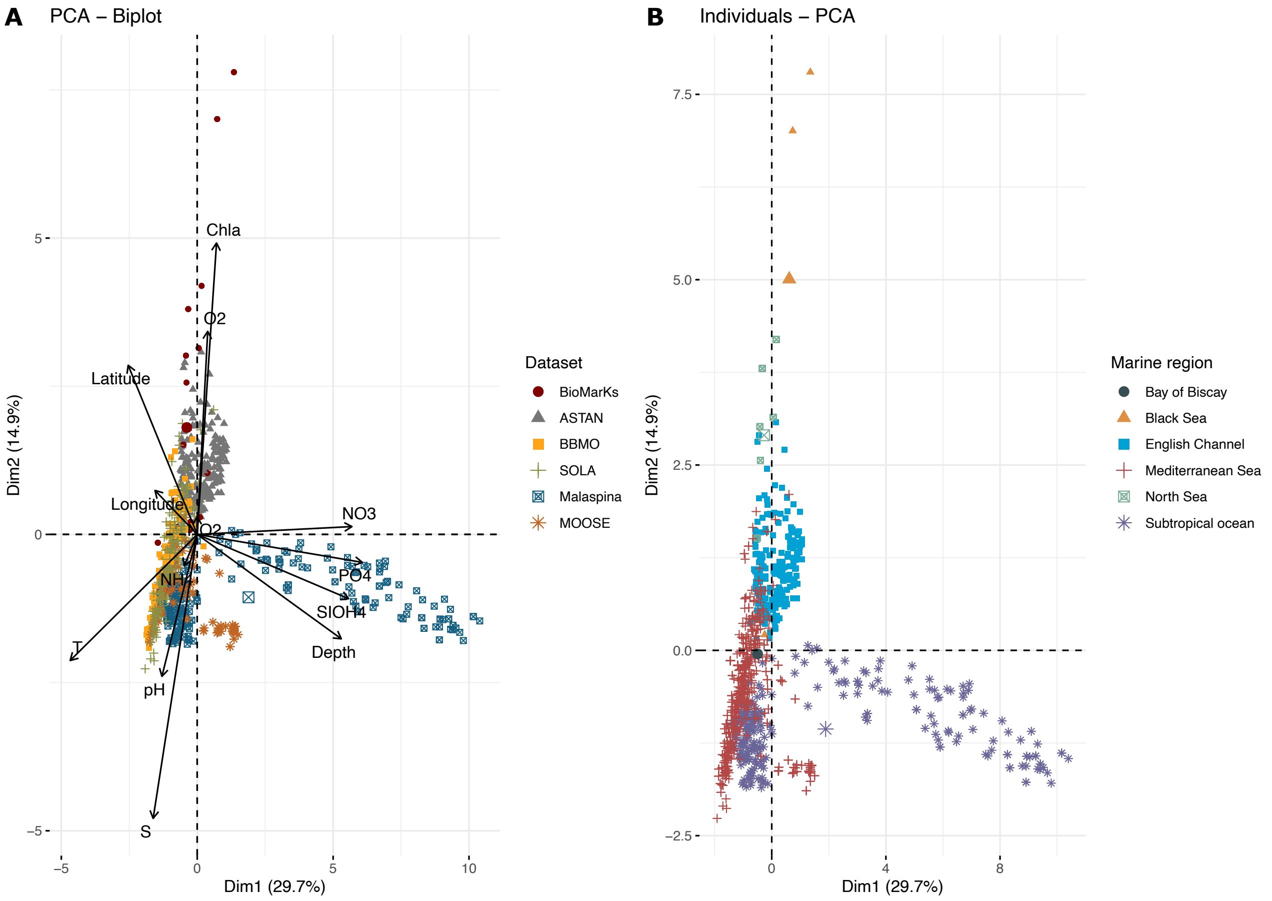

**Fig. S3:** Relative number of unassigned sequences across the taxonomic hierarchy (Kingdom to Species) for each dataset. Colors represent the 6 studied datasets. (A) Relative number of unassigned sequences across the homogenisation workflow: initial taxonomy of metabarcodes upon dataset receival (top); after taxonomic assignment with PR2 database v4.12.0 (middle; Fig. S1 (A) step 2); after filtering sequences by length and contamination check (bottom; Fig. S1 (A) step 3 & 4). (B) Relative number (top) and relative proportion (bottom) of unassigned sequences after the homogenisation workflow.

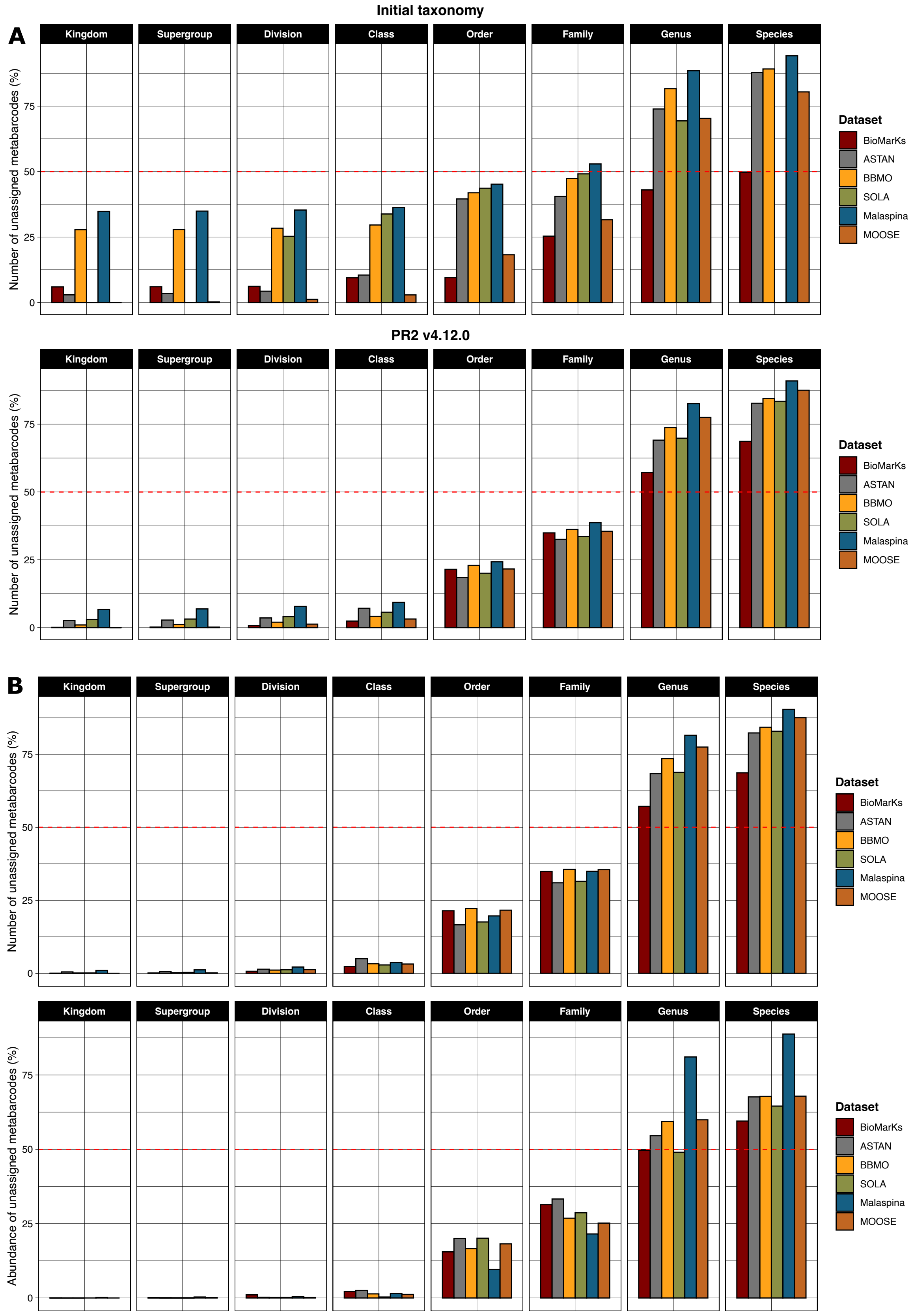

**Fig. S4:** Distribution of unassigned metabarcodes at kingdom level. (A) Map indicating the sampling sites of samples containing the studied metabarcodes. (B) Number of metabarcodes found across each dataset. 89.6% of metabarcodes are found in the Malaspina dataset. (C) Number of metabarcodes found across each depth type within the Malaspina dataset. The metabarcodes found in the bathypelagic layer represent 87.7% of metabarcodes unassigned at kingdom level.

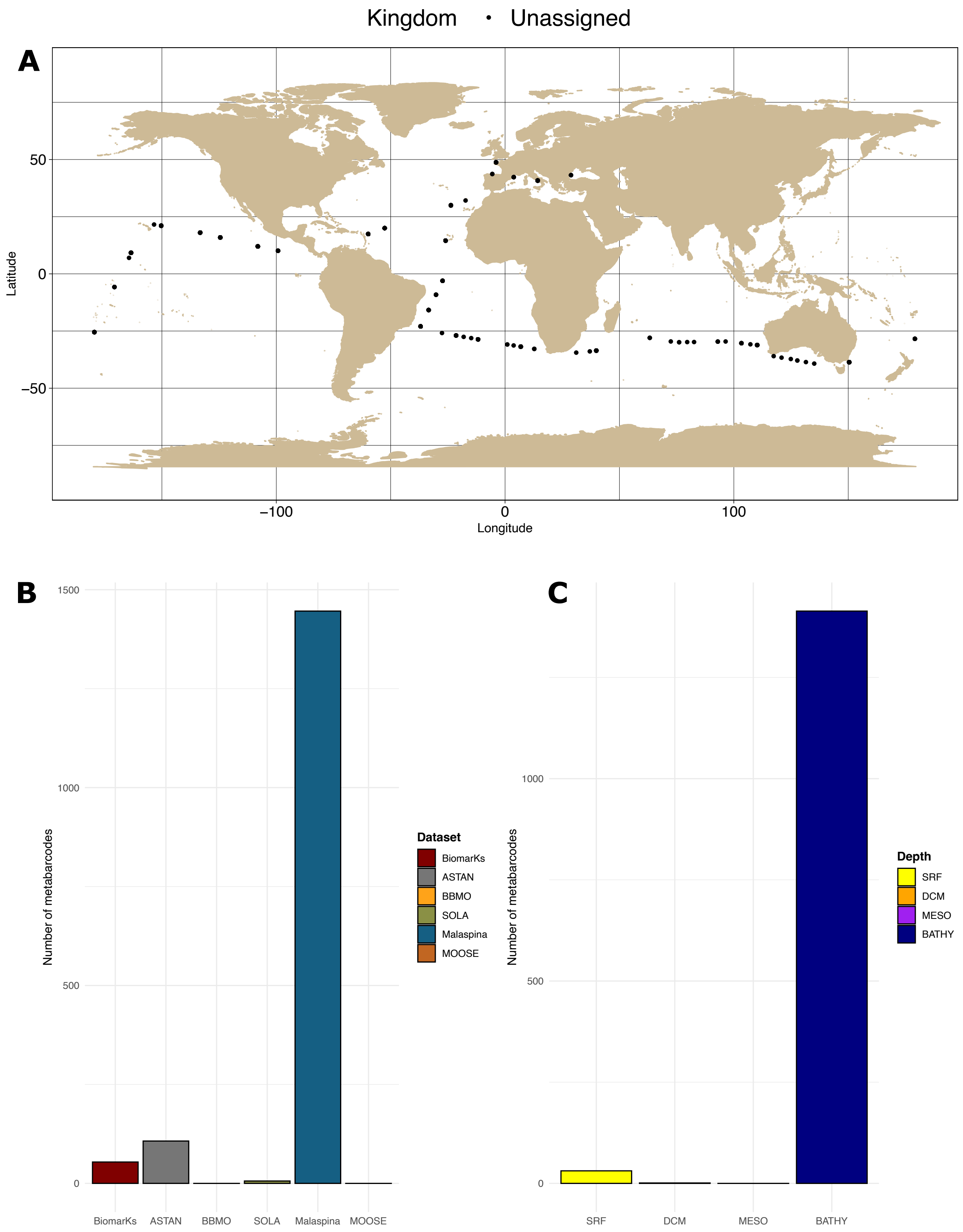

**Fig. S5:** A-diversity and abundance of metabarcodes lacking taxonomic annotation at the genus level, across datasets and major protist divisions. (A) Relative number (top) and relative abundance (bottom) of unassigned metabarcodes. Unassigned sequences are grouped according to their taxonomic annotation at the level of the Division (indicated by the color code). Only the top 4 most diverse / abundant divisions are plotted. The rest of divisions are included in “Others” and represent at no more than 6% of metabarcodes each. (B) Shannon diversity index of top protist divisions (in terms of diversity and abundance) for each dataset. The index was calculated taking into account both assigned and unassigned sequences (top), only assigned sequences (middle) and only unassigned sequences (bottom).

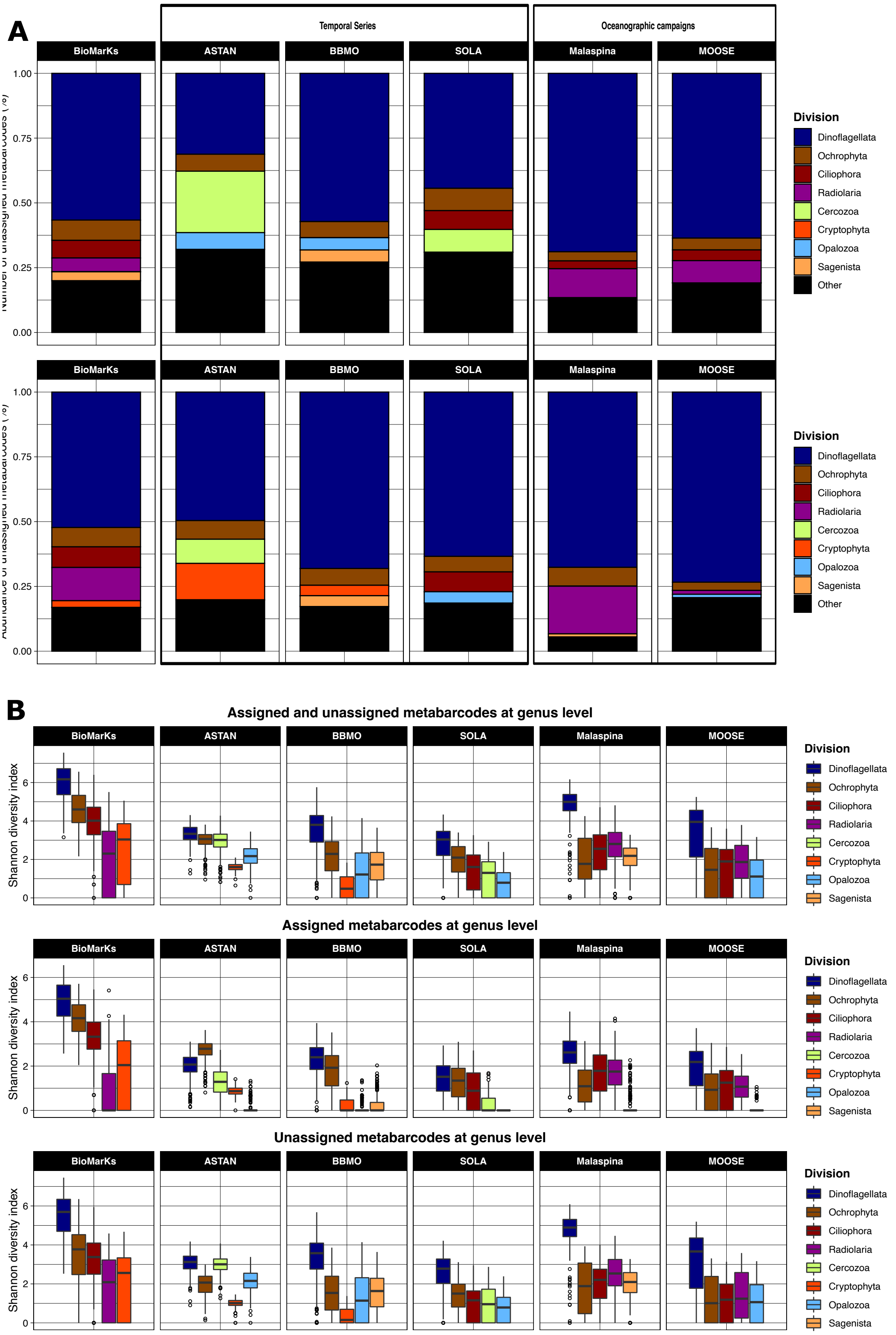

**Fig. S6:** Representativity of Dinoflagellata Classes in the 6 metabarcoding datasets. (A) Relative number (top) and relative abundance (bottom) of Dinoflagellata Classes. Others include Classes representing on average < 1% of Dinoflagellata metabarcodes in terms of number (n=3 655) and <1% in terms of abundance. These Classes are: Noctilucophyceae (3 287 metabarcodes), Oxyrrhea (1 metabarcode), Ellobiophyceae (12 metabarcodes) and sequences of Unknown class (no taxonomic assignment at Class level, 355 metabarcodes). (B) Relative number (top) and relative abundance (bottom) of assigned and unassigned sequences among Dinophyceae (left) and Syndiniales (right).

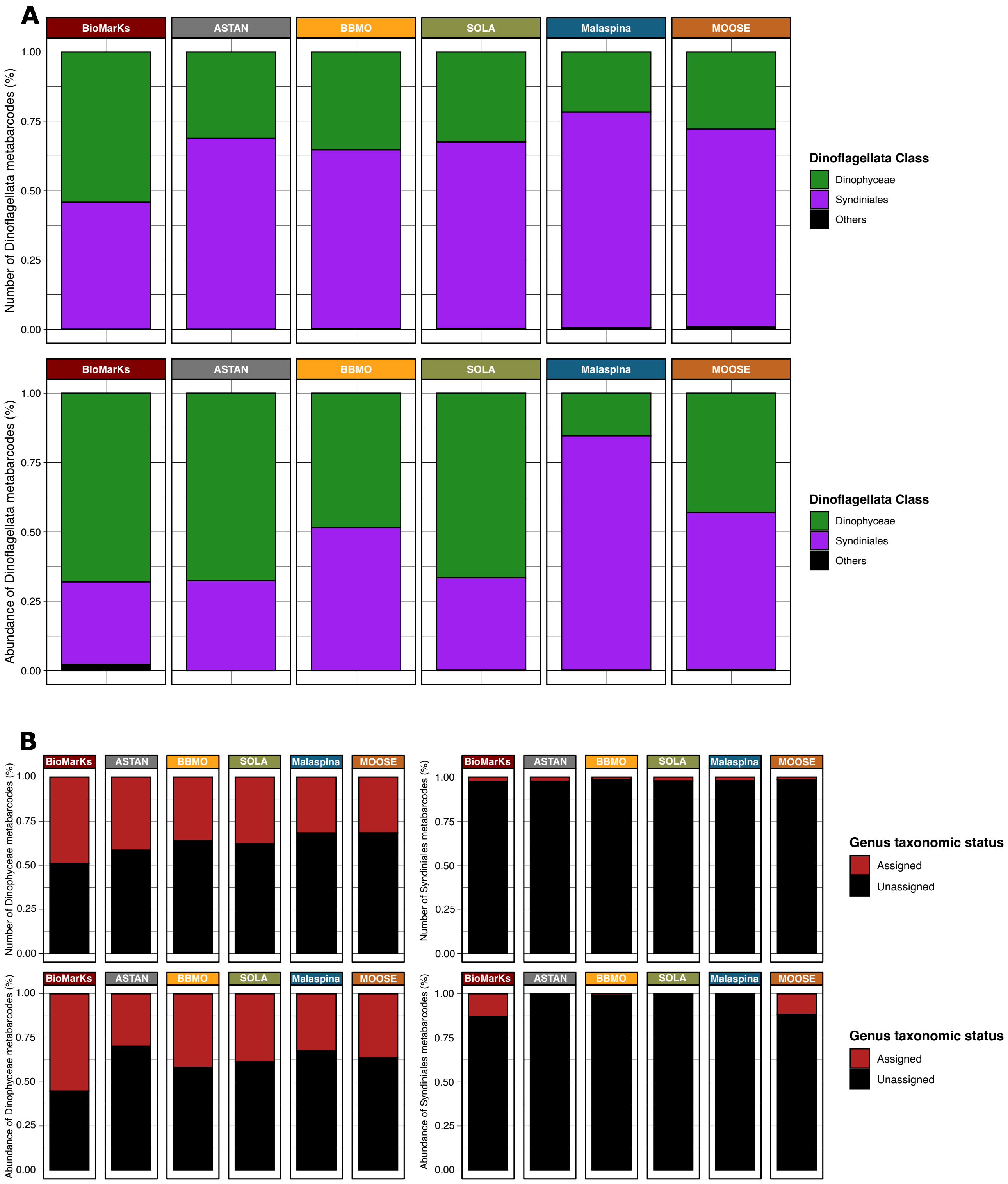

**Fig. S7:** Syndiniales CCs size and taxonomy. (A) Size distribution of Syndiniales according to their assignment at genus level: unassigned CCs (4 245 CCs, top) and assigned CCs (72 CCs, bottom). Scale bars differ between the two graphs. (B) Number of unassigned Syndiniales Connected Components (CCs) in the network (12 619 CCs) across Syndiniales orders. A CC is considered unassigned if it is composed of metabarcodes lacking taxonomic affiliation at the genus level. The network contained both assigned and unassigned CCs. The total number of Syndiniales CCs in the network is 4 317 (34.21% of the network) and the number of unassigned Syndiniales CCs is 4 245 (33.64% of the network; 47.60% of unassigned CCs of the network (i.e. 8 919 CCs); 98.33% of Syndiniales CCs). Syndiniales order IV (Dino-Group-IV) was composed of 41 CCs which contained only assigned sequences (genera *Hematodinium* and *Syndinium*) and was, thus, not plotted. "Unknown" corresponds to CCs composed of metabarcodes lacking taxonomic affiliation both at the order level.

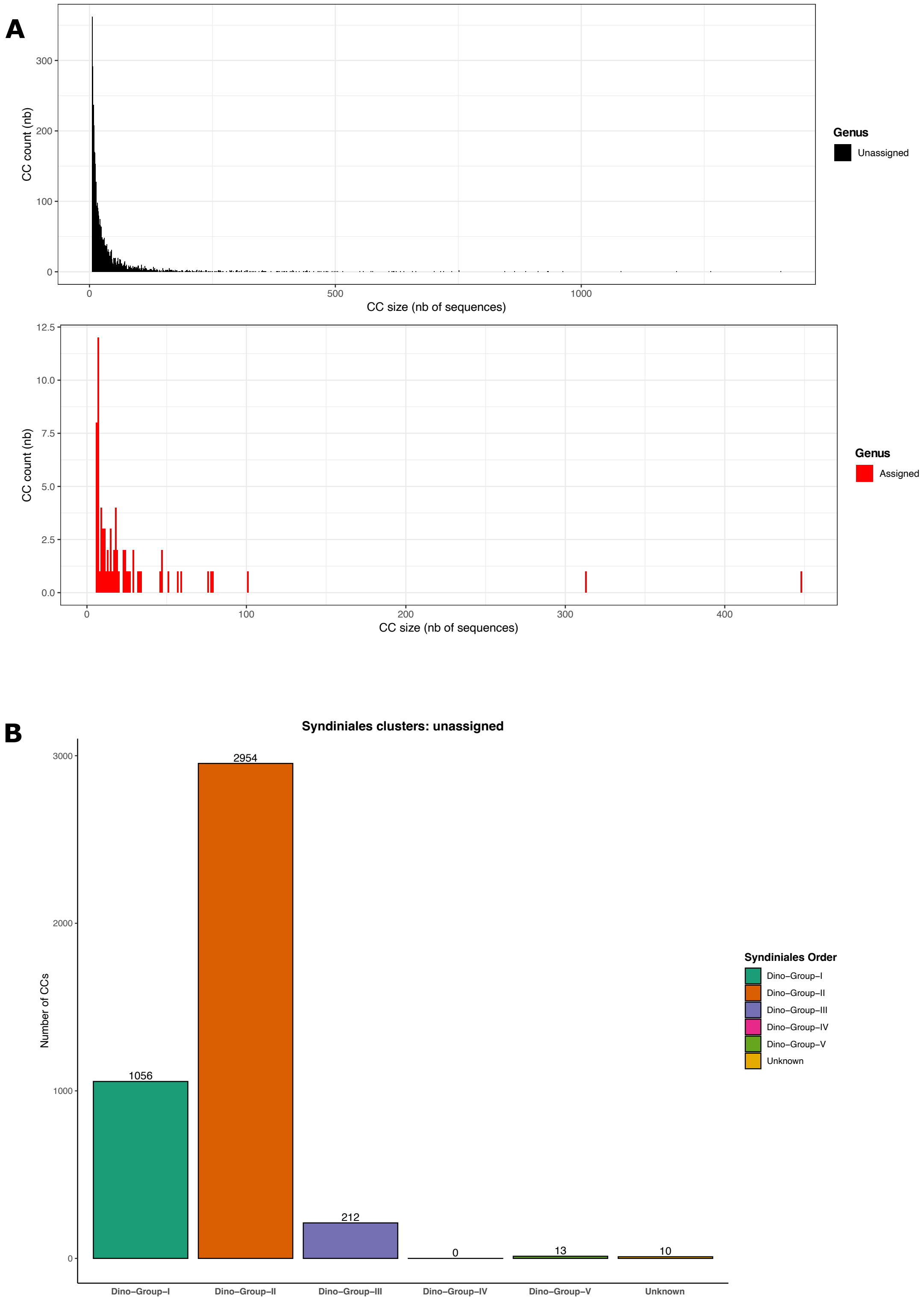

**Fig. S8:** Relative abundance of Syndiniales CCs across the 3 time-series: (A) ASTAN, (B) BBMO, (C) SOLA. The average monthly abundance is calculated per year and compiled as an average of the dataset. Colors indicate the presence/absence of taxonomic assignment of the CC at the genus level (red: presence of taxonomy; known genus and gray: absence of taxonomy; unassigned/unknown genus). Scale bars differ between the graphs. The big dots represent the CCs that were selected by the Escoufier's equivalent vectors and Lomb-Scargle periodogram algorithm methods for each time-series.

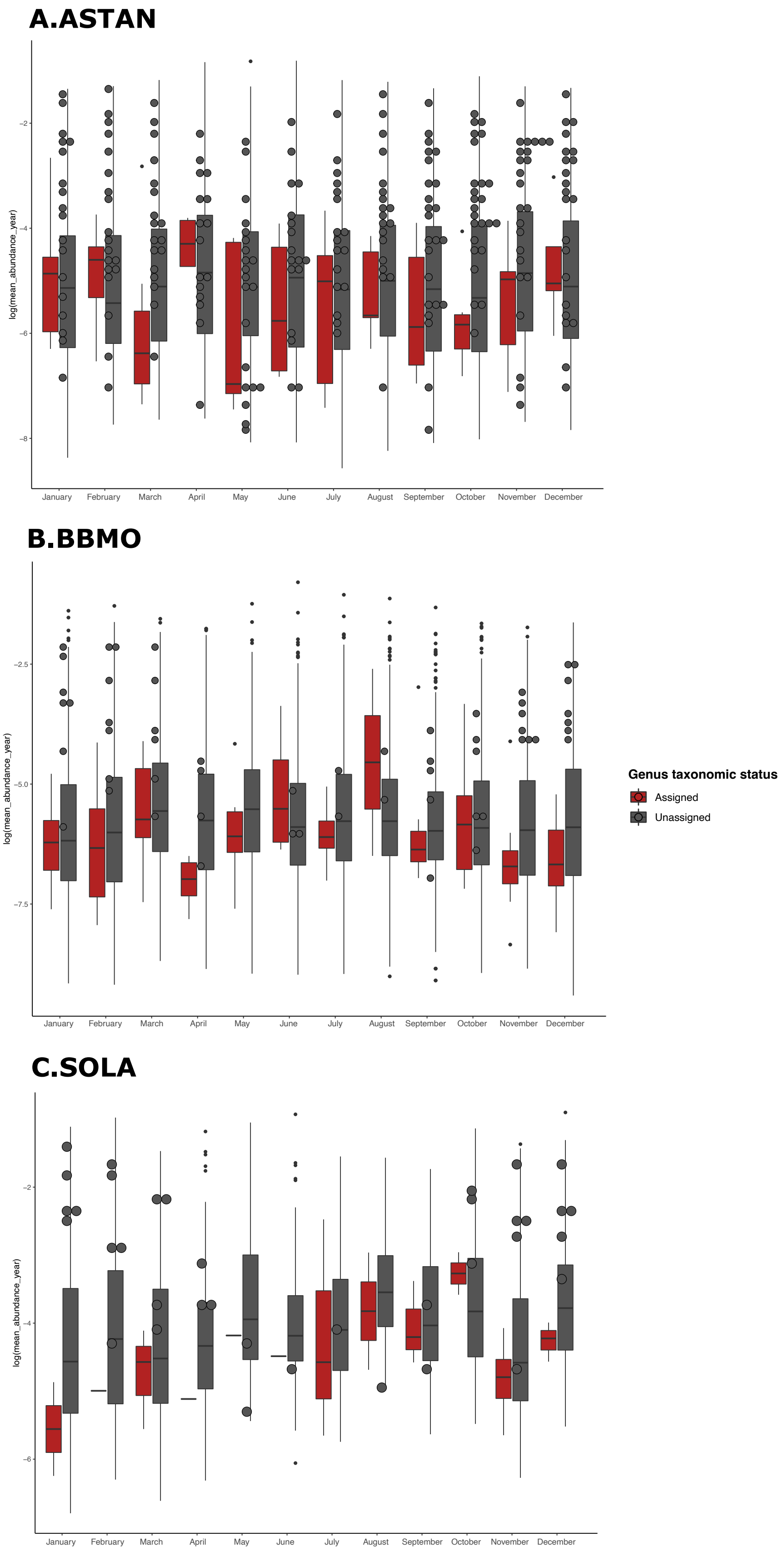

### Supplementary Tables

**Table S1:** Information about the 6 metabarcoding datasets included in this study.

| | BioMarKs | ASTAN | BBMO | SOLA | Malaspina | MOOSE | Global dataset |
|---|---|---|---|---|---|---|---|
| **Provider** | R Logares | M Caracciolo | R Logares | P Galand | R Logares | M Mendez-Sandin | - |
| **Abundance table** | Provided | Provided | Provided | Provided | Provided | Provided | - |
| **Taxonomic affiliation** | Provided | Provided | Provided | Provided | Provided | Provided | - |
| **DB version of starting taxonomy** | NA | PR2 v4.12.0 | PR2 v4.11.1 | SILVA v128 | PR2 v4.11.1 | PR2 v4.12.0 | - |
| **Metadata origin** | SOMLIT, World Ocean Database, SeaDataNet | SOMLIT | Provided | Provided | Provided | Provided | - |
| **Samples (nb)** | 115 | 374 | 327 | 154 | 289 | 272 | 1531 |
| **Stations (nb)** | 6 | 1 | 1 | 1 | 122 | 26 | 155 |
| **Vertical profile max depth** | 100 | 3 | 3 | 3 | 4000 | 2000 | - |
| **Different months** | 6 | 12 | 12 | 12 | 8 | 3 | - |
| **Years** | 1 | 8 | 10 | 8 | 1 | 2 | - |
| **Sequencing** | 454 | Illumina MiSeq | Illumina MiSeq | Illumina MiSeq | Illumina MiSeq | Illumina MiSeq | - |
| **Clusters of reads** | OTU | ASV | ASV | ASV | ASV | ASV | - |
| **Read clustering threshold** | NA | NA | 95 % | NA | 95 % | 80 % | - |
| **Applied read clustering threshold in this study** | 80 % | 80 % | - | 80 % | - | 80 % | - |
| **Starting number of metabarcodes** | 423547 | 7933 | 59428 | 6400 | 59428 | 12279 | 569015 |
| **Filtering step 1 (id > 80%, only protist sequences kept)** | 239508 | 6239 | 29682 | 5554 | 59423 | 11322 | 351728 |
| **Filtering step 2: taxonomic assignment specifications** | PR2 v4.12.0, eval 0.01 | PR2 v4.12.0, eval 0.01 | PR2 v4.12.0, eval 0.01 | PR2 v4.12.0, eval 0.01 | PR2 v4.12.0, eval 0.01 | PR2 v4.12.0, eval 0.01 | PR2 v4.12.0, eval 0.01 |
| **Multicellular lineages removed** | Metazoa, Streptophyta, Florideophyceae, Bangiophyceae, Phaeophyceae, Ulvophyceae | Metazoa, Streptophyta, Florideophyceae, Bangiophyceae, Phaeophyceae, Ulvophyceae | Metazoa, Streptophyta, Florideophyceae, Bangiophyceae, Phaeophyceae, Ulvophyceae | Metazoa, Streptophyta, Florideophyceae, Bangiophyceae, Phaeophyceae, Ulvophyceae | Metazoa, Streptophyta, Florideophyceae, Bangiophyceae, Phaeophyceae, Ulvophyceae | Metazoa, Streptophyta, Florideophyceae, Bangiophyceae, Phaeophyceae, Ulvophyceae | Metazoa, Streptophyta, Florideophyceae, Bangiophyceae, Phaeophyceae, Ulvophyceae |
| **Filtering step 3 (sequence length > 200 bp)** | 236618 | 6213 | 29073 | 5276 | 58555 | 11304 | 347039 |
| **Filtering step 4 (contamination check with SILVA v138)** | 236575 | 6087 | 28809 | 5218 | 55173 | 11303 | 343165 |

**Table S2:** Jaccard diversity index calculated between sea regions for the following protist groups: Dinophyeae, Ciliophora, Radiolaria, Ochrophyta, Cercozoa, Cryptophyta, Opalozoa, Sagenista and Syndiniales.

| **Dinophyceae** | **Bay.of.Biscay** | **Black.Sea** | **English.Channel** | **Mediterranean.Sea** | **North.Sea** | **Subtropical.ocean** |
|---|---|---|---|---|---|---|
| **Bay of Biscay** | 0 | 0.938173454130847 | 0.87497104538359 | 0.804672833794989 | 0.927670261180023 | 0.818861390415376 |
| **Black Sea** | 0.938173454130847 | 0 | 0.901578438140029 | 0.890869330127625 | 0.791834766178996 | 0.962644200953342 |
| **English Channel** | 0.87497104538359 | 0.901578438140029 | 0 | 0.798497443157509 | 0.859471772489166 | 0.894607381909587 |
| **Mediterranean Sea** | 0.804672833794989 | 0.890869330127625 | 0.798497443157509 | 0 | 0.867745884222841 | 0.664604534857357 |
| **North Sea** | 0.927670261180023 | 0.791834766178996 | 0.859471772489166 | 0.867745884222841 | 0 | 0.954859723071374 |
| **Subtropical ocean** | 0.818861390415376 | 0.962644200953342 | 0.894607381909587 | 0.664604534857357 | 0.954859723071374 | 0 |
| **Ciliophora** | Bay.of.Biscay | Black.Sea | English.Channel | Mediterranean.Sea | North.Sea | Subtropical.ocean |
| **Bay of Biscay** | 0 | 0.867381892806053 | 0.902947145619635 | 0.842217097011825 | 0.872886640938259 | 0.858727477402051 |
| **Black Sea** | 0.867381892806053 | 0 | 0.887898854315709 | 0.891501486515274 | 0.79049701022653 | 0.944252209613938 |
| **English Channel** | 0.902947145619635 | 0.887898854315709 | 0 | 0.777126046175384 | 0.803651192794572 | 0.923737512275255 |
| **Mediterranean Sea** | 0.842217097011825 | 0.891501486515274 | 0.777126046175384 | 0 | 0.828880954082161 | 0.729483694176171 |
| **North Sea** | 0.872886640938259 | 0.79049701022653 | 0.803651192794572 | 0.828880954082161 | 0 | 0.92877157288191 |
| **Subtropical ocean** | 0.858727477402051 | 0.944252209613938 | 0.923737512275255 | 0.729483694176171 | 0.92877157288191 | 0 |
| **Radiolaria** | Bay.of.Biscay | Black.Sea | English.Channel | Mediterranean.Sea | North.Sea | Subtropical.ocean |
| **Bay of Biscay** | 0 | 0.97462797037596 | 0.870508797970726 | 0.942580875095232 | 0.914028700372153 | 0.98111749754938 |
| **Black Sea** | 0.97462797037596 | 0 | 0.958390416328276 | 0.990831276360653 | 0.983866226229662 | 0.991593478316324 |
| **English Channel** | 0.870508797970726 | 0.958390416328276 | 0 | 0.961150318767074 | 0.899296029993727 | 0.986294136252654 |
| **Mediterranean Sea** | 0.942580875095232 | 0.990831276360653 | 0.961150318767074 | 0 | 0.896220133271001 | 0.727417292423973 |
| **North Sea** | 0.914028700372153 | 0.983866226229662 | 0.899296029993727 | 0.896220133271001 | 0 | 0.993423735613338 |
| **Subtropical ocean** | 0.98111749754938 | 0.991593478316324 | 0.986294136252654 | 0.727417292423973 | 0.993423735613338 | 0 |
| **Ochrophyta** | Bay.of.Biscay | Black.Sea | English.Channel | Mediterranean.Sea | North.Sea | Subtropical.ocean |
| **Bay of Biscay** | 0 | 0.944213049494936 | 0.953554574430131 | 0.862135906227205 | 0.939353323258805 | 0.839757547138781 |
| **Black Sea** | 0.944213049494936 | 0 | 0.918409472383869 | 0.892072704102568 | 0.732692288501858 | 0.961403091261643 |
| **English Channel** | 0.953554574430131 | 0.918409472383869 | 0 | 0.806755268471589 | 0.882128402710598 | 0.953440648813368 |
| **Mediterranean Sea** | 0.862135906227205 | 0.892072704102568 | 0.806755268471589 | 0 | 0.83361488993896 | 0.834034881022253 |
| **North Sea** | 0.939353323258805 | 0.732692288501858 | 0.882128402710598 | 0.83361488993896 | 0 | 0.965028633968432 |
| **Subtropical ocean** | 0.839757547138781 | 0.961403091261643 | 0.953440648813368 | 0.834034881022253 | 0.965028633968432 | 0 |
| **Cercozoa** | Bay.of.Biscay | Black.Sea | English.Channel | Mediterranean.Sea | North.Sea | Subtropical.ocean |
| **Bay of Biscay** | 0 | 0.982103174330624 | 0.96701345562615 | 0.878828971450338 | 0.904605092480974 | 0.930785824223574 |
| **Black Sea** | 0.982103174330624 | 0 | 0.951501456544751 | 0.927806049444131 | 0.877524702758534 | 0.99158348108881 |
| **English Channel** | 0.96701345562615 | 0.951501456544751 | 0 | 0.745475896500063 | 0.843165738440751 | 0.967596506393834 |
| **Mediterranean Sea** | 0.878828971450338 | 0.927806049444131 | 0.745475896500063 | 0 | 0.837231980758612 | 0.88421222368964 |
| **North Sea** | 0.904605092480974 | 0.877524702758534 | 0.843165738440751 | 0.837231980758612 | 0 | 0.981476463546255 |
| **Subtropical ocean** | 0.930785824223574 | 0.99158348108881 | 0.967596506393834 | 0.88421222368964 | 0.981476463546255 | 0 |
| **Cryptophyta** | Bay.of.Biscay | Black.Sea | English.Channel | Mediterranean.Sea | North.Sea | Subtropical.ocean |
| **Bay of Biscay** | 0 | 0.852266116053246 | 0.881612974786431 | 0.747806749032386 | 0.944704795765131 | 0.791666798740924 |
| **Black Sea** | 0.852266116053246 | 0 | 0.821118584548414 | 0.735923947815264 | 0.671563329597875 | 0.928054816498048 |
| **English Channel** | 0.881612974786431 | 0.821118584548414 | 0 | 0.786600033491997 | 0.813523218679703 | 0.870377288638509 |
| **Mediterranean Sea** | 0.747806749032386 | 0.735923947815264 | 0.786600033491997 | 0 | 0.850928644235467 | 0.882593230983105 |
| **North Sea** | 0.944704795765131 | 0.671563329597875 | 0.813523218679703 | 0.850928644235467 | 0 | 0.938662701091859 |
| **Subtropical ocean** | 0.791666798740924 | 0.928054816498048 | 0.870377288638509 | 0.882593230983105 | 0.938662701091859 | 0 |
| **Opalozoa** | Bay.of.Biscay | Black.Sea | English.Channel | Mediterranean.Sea | North.Sea | Subtropical.ocean |
| **Bay of Biscay** | 0 | 1 | 0.900792047825135 | 0.810741256648265 | 0.915114872445857 | 0.904576802623282 |
| **Black Sea** | 1 | 0 | 0.924648613090138 | 0.976234364064496 | 0.874080566513386 | 0.971848215239348 |
| **English Channel** | 0.900792047825135 | 0.924648613090138 | 0 | 0.745279921844327 | 0.770950207818785 | 0.90305636237645 |
| **Mediterranean Sea** | 0.810741256648265 | 0.976234364064496 | 0.745279921844327 | 0 | 0.839940179196507 | 0.843786084383779 |

| | | | | | | |
|---|---|---|---|---|---|---|
| **North Sea** | 0.915114872445857 | 0.874080566513386 | 0.770950207818785 | 0.839940179196507 | 0 | 0.957003482122313 |
| **Subtropical ocean** | 0.904576802623282 | 0.971848215239348 | 0.90305636237645 | 0.843786084383779 | 0.957003482122313 | 0 |
| **Sagenista** | Bay.of.Biscay | Black.Sea | English.Channel | Mediterranean.Sea | North.Sea | Subtropical.ocean |
| **Bay of Biscay** | 0 | 0.927494765810393 | 0.812106830102491 | 0.746219107052659 | 0.847799688731939 | 0.760141643941602 |
| **Black Sea** | 0.927494765810393 | 0 | 0.83502956899843 | 0.927283159174829 | 0.704648885459417 | 0.970443781190252 |
| **English Channel** | 0.812106830102491 | 0.83502956899843 | 0 | 0.738032807799124 | 0.682014630383829 | 0.845932127615074 |
| **Mediterranean Sea** | 0.746219107052659 | 0.927283159174829 | 0.738032807799124 | 0 | 0.809437286678021 | 0.620999021505498 |
| **North Sea** | 0.847799688731939 | 0.704648885459417 | 0.682014630383829 | 0.809437286678021 | 0 | 0.904155003240767 |
| **Subtropical ocean** | 0.760141643941602 | 0.970443781190252 | 0.845932127615074 | 0.620999021505498 | 0.904155003240767 | 0 |
| **Syndiniales** | Bay.of.Biscay | Black.Sea | English.Channel | Mediterranean.Sea | North.Sea | Subtropical.ocean |
| **Bay of Biscay** | 0 | 0.940557162491321 | 0.872991324662993 | 0.867376931604074 | 0.910315399979933 | 0.898494917470853 |
| **Black Sea** | 0.940557162491321 | 0 | 0.91577064018812 | 0.949739577024946 | 0.817131780391512 | 0.96862939955675 |
| **English Channel** | 0.872991324662993 | 0.91577064018812 | 0 | 0.814610404707088 | 0.84839192562793 | 0.872386318763916 |
| **Mediterranean Sea** | 0.867376931604074 | 0.949739577024946 | 0.814610404707088 | 0 | 0.902815004053065 | 0.60089678886582 |
| **North Sea** | 0.910315399979933 | 0.817131780391512 | 0.84839192562793 | 0.902815004053065 | 0 | 0.935820962510432 |
| **Subtropical ocean** | 0.898494917470853 | 0.96862939955675 | 0.872386318763916 | 0.60089678886582 | 0.935820962510432 | 0 |

**Table S3:** Indicator CCs indicated by the Escouffier's equivalent method.

| | Dataset | CC_id | Escouffier | RV |
|---|---|---|---|---|
| **1** | ASTAN | CC_unknown_535 | 1 | 0.248 |
| **2** | ASTAN | CC_unknown_154 | 2 | 0.32 |
| **3** | ASTAN | CC_unknown_532 | 3 | 0.372 |
| **4** | ASTAN | CC_unknown_168 | 4 | 0.409 |
| **5** | ASTAN | CC_unknown_547 | 5 | 0.442 |
| **6** | ASTAN | CC_unknown_368 | 6 | 0.471 |
| **7** | ASTAN | CC_unknown_13 | 7 | 0.491 |
| **8** | ASTAN | CC_unknown_109 | 8 | 0.511 |
| **9** | ASTAN | CC_unknown_363 | 9 | 0.53 |
| **10** | ASTAN | CC_unknown_257 | 10 | 0.544 |
| **11** | ASTAN | CC_unknown_498 | 11 | 0.557 |
| **12** | ASTAN | CC_unknown_370 | 12 | 0.57 |
| **13** | ASTAN | CC_unknown_244 | 13 | 0.581 |
| **14** | ASTAN | CC_unknown_172 | 14 | 0.592 |
| **15** | ASTAN | CC_unknown_77 | 15 | 0.602 |
| **16** | ASTAN | CC_unknown_210 | 16 | 0.611 |
| **17** | ASTAN | CC_known_12 | 17 | 0.619 |
| **18** | ASTAN | CC_unknown_183 | 18 | 0.627 |
| **19** | ASTAN | CC_unknown_550 | 19 | 0.634 |
| **20** | ASTAN | CC_unknown_35 | 20 | 0.641 |
| **21** | ASTAN | CC_unknown_30 | 21 | 0.648 |
| **22** | ASTAN | CC_unknown_254 | 22 | 0.655 |
| **23** | ASTAN | CC_unknown_80 | 23 | 0.661 |
| **24** | ASTAN | CC_unknown_307 | 24 | 0.667 |
| **25** | ASTAN | CC_unknown_441 | 25 | 0.672 |
| **26** | ASTAN | CC_unknown_462 | 26 | 0.678 |
| **27** | ASTAN | CC_unknown_530 | 27 | 0.683 |
| **28** | ASTAN | CC_unknown_126 | 28 | 0.689 |
| **29** | ASTAN | CC_unknown_328 | 29 | 0.693 |
| **30** | ASTAN | CC_unknown_209 | 30 | 0.697 |
| **31** | ASTAN | CC_unknown_134 | 31 | 0.702 |
| **32** | ASTAN | CC_unknown_339 | 32 | 0.706 |
| **33** | ASTAN | CC_unknown_272 | 33 | 0.71 |
| **34** | ASTAN | CC_unknown_496 | 34 | 0.714 |
| **35** | ASTAN | CC_unknown_295 | 35 | 0.717 |
| **36** | ASTAN | CC_unknown_483 | 36 | 0.721 |
| **37** | ASTAN | CC_unknown_553 | 37 | 0.725 |
| **38** | ASTAN | CC_unknown_20 | 38 | 0.728 |
| **39** | ASTAN | CC_unknown_12 | 39 | 0.731 |
| **40** | ASTAN | CC_unknown_349 | 40 | 0.734 |
| **41** | ASTAN | CC_unknown_58 | 41 | 0.737 |
| **42** | ASTAN | CC_unknown_62 | 42 | 0.74 |
| **43** | ASTAN | CC_unknown_362 | 43 | 0.743 |
| **44** | ASTAN | CC_unknown_227 | 44 | 0.746 |
| **45** | ASTAN | CC_unknown_320 | 45 | 0.749 |
| **46** | BBMO | CC_unknown_72 | 1 | 0.261 |
| **47** | BBMO | CC_unknown_330 | 2 | 0.344 |
| **48** | BBMO | CC_unknown_112 | 3 | 0.403 |
| **49** | BBMO | CC_unknown_562 | 4 | 0.442 |
| **50** | BBMO | CC_unknown_213 | 5 | 0.478 |
| **51** | BBMO | CC_unknown_2182 | 6 | 0.507 |
| **52** | BBMO | CC_unknown_1964 | 7 | 0.532 |
| **53** | BBMO | CC_unknown_401 | 8 | 0.552 |
| **54** | BBMO | CC_unknown_1225 | 9 | 0.567 |
| **55** | BBMO | CC_unknown_926 | 10 | 0.58 |
| **56** | BBMO | CC_unknown_129 | 11 | 0.592 |
| **57** | BBMO | CC_unknown_389 | 12 | 0.604 |
| **58** | BBMO | CC_unknown_154 | 13 | 0.615 |
| **59** | BBMO | CC_unknown_767 | 14 | 0.626 |
| **60** | BBMO | CC_unknown_1451 | 15 | 0.636 |
| **61** | BBMO | CC_unknown_1869 | 16 | 0.645 |
| **62** | BBMO | CC_unknown_1827 | 17 | 0.653 |
| **63** | BBMO | CC_unknown_1514 | 18 | 0.661 |
| **64** | BBMO | CC_unknown_1868 | 19 | 0.668 |
| **65** | BBMO | CC_unknown_1039 | 20 | 0.675 |
| **66** | BBMO | CC_unknown_1532 | 21 | 0.68 |

| | | | | |
|---|---|---|---|---|
| **67** | BBMO | CC_unknown_482 | 22 | 0.686 |
| **68** | BBMO | CC_unknown_2139 | 23 | 0.691 |
| **69** | BBMO | CC_unknown_404 | 24 | 0.696 |
| **70** | BBMO | CC_unknown_989 | 25 | 0.701 |
| **71** | BBMO | CC_unknown_604 | 26 | 0.706 |
| **72** | BBMO | CC_unknown_161 | 27 | 0.711 |
| **73** | BBMO | CC_unknown_1960 | 28 | 0.716 |
| **74** | BBMO | CC_unknown_2212 | 29 | 0.721 |
| **75** | BBMO | CC_unknown_839 | 30 | 0.725 |
| **76** | BBMO | CC_unknown_894 | 31 | 0.729 |
| **77** | BBMO | CC_unknown_1775 | 32 | 0.733 |
| **78** | BBMO | CC_unknown_1893 | 33 | 0.737 |
| **79** | BBMO | CC_unknown_296 | 34 | 0.74 |
| **80** | BBMO | CC_unknown_1563 | 35 | 0.744 |
| **81** | BBMO | CC_unknown_126 | 36 | 0.747 |
| **82** | SOLA | CC_unknown_425 | 1 | 0.341 |
| **83** | SOLA | CC_unknown_164 | 2 | 0.436 |
| **84** | SOLA | CC_unknown_918 | 3 | 0.512 |
| **85** | SOLA | CC_unknown_126 | 4 | 0.551 |
| **86** | SOLA | CC_unknown_199 | 5 | 0.579 |
| **87** | SOLA | CC_unknown_149 | 6 | 0.604 |
| **88** | SOLA | CC_unknown_326 | 7 | 0.625 |
| **89** | SOLA | CC_unknown_39 | 8 | 0.643 |
| **90** | SOLA | CC_unknown_892 | 9 | 0.658 |
| **91** | SOLA | CC_unknown_171 | 10 | 0.673 |
| **92** | SOLA | CC_unknown_226 | 11 | 0.686 |
| **93** | SOLA | CC_unknown_1793 | 12 | 0.697 |
| **94** | SOLA | CC_unknown_143 | 13 | 0.707 |
| **95** | SOLA | CC_unknown_183 | 14 | 0.717 |
| **96** | SOLA | CC_unknown_333 | 15 | 0.728 |
| **97** | SOLA | CC_unknown_293 | 16 | 0.737 |
| **98** | SOLA | CC_unknown_1908 | 17 | 0.747 |

**Table S4:** Rhythmic CCs indicated by the Lomb-Scargle Periodogram (LSP) method.

| | Dataset | CC_id | PNmax | Pvalue | Period |
|---|---|---|---|---|---|
| 1 | ASTAN | CC_known_1 | 14.99 | 1E-04 | 363.25 |
| 2 | ASTAN | CC_known_14 | 38.403 | 0 | 372.56 |
| 3 | ASTAN | CC_known_2 | 13.533 | 5E-04 | 854.71 |
| 4 | ASTAN | CC_known_4 | 10.914 | 0.0068 | 372.56 |
| 5 | ASTAN | CC_unknown_1 | 15.19 | 1E-04 | 392.7 |
| 6 | ASTAN | CC_unknown_10 | 16.506 | 0 | 372.56 |
| 7 | ASTAN | CC_unknown_100 | 28.65 | 0 | 372.56 |
| 8 | ASTAN | CC_unknown_103 | 34.521 | 0 | 372.56 |
| 9 | ASTAN | CC_unknown_104 | 11.247 | 0.0049 | 363.25 |
| 10 | ASTAN | CC_unknown_106 | 15.962 | 0 | 363.25 |
| 11 | ASTAN | CC_unknown_107 | 10.778 | 0.0078 | 363.25 |
| 12 | ASTAN | CC_unknown_11 | 27.452 | 0 | 363.25 |
| 13 | ASTAN | CC_unknown_110 | 10.851 | 0.0072 | 454.06 |
| 14 | ASTAN | CC_unknown_111 | 11.814 | 0.0028 | 2906 |
| 15 | ASTAN | CC_unknown_112 | 11.592 | 0.0034 | 345.95 |
| 16 | ASTAN | CC_unknown_113 | 10.096 | 0.0152 | 363.25 |
| 17 | ASTAN | CC_unknown_116 | 13.206 | 7E-04 | 2421.7 |
| 18 | ASTAN | CC_unknown_118 | 14.369 | 2E-04 | 363.25 |
| 19 | ASTAN | CC_unknown_119 | 59.04 | 0 | 372.56 |
| 20 | ASTAN | CC_unknown_12 | 13.212 | 7E-04 | 354.39 |
| 21 | ASTAN | CC_unknown_120 | 16.056 | 0 | 726.5 |
| 22 | ASTAN | CC_unknown_121 | 10.003 | 0.0167 | 363.25 |
| 23 | ASTAN | CC_unknown_122 | 10.615 | 0.0091 | 345.95 |
| 24 | ASTAN | CC_unknown_123 | 10.134 | 0.0147 | 220.15 |
| 25 | ASTAN | CC_unknown_125 | 11.599 | 0.0034 | 372.56 |
| 26 | ASTAN | CC_unknown_126 | 30.339 | 0 | 2906 |
| 27 | ASTAN | CC_unknown_128 | 38.644 | 0 | 363.25 |
| 28 | ASTAN | CC_unknown_129 | 18.939 | 0 | 372.56 |
| 29 | ASTAN | CC_unknown_13 | 102.62 | 0 | 363.25 |
| 30 | ASTAN | CC_unknown_131 | 10.42 | 0.011 | 345.95 |
| 31 | ASTAN | CC_unknown_132 | 32.41 | 0 | 363.25 |
| 32 | ASTAN | CC_unknown_134 | 47.33 | 0 | 363.25 |

| | | | | | |
|---|---|---|---|---|---|
| 33 | ASTAN | CC_unknown_135 | 14.175 | 3E-04 | 372.56 |
| 34 | ASTAN | CC_unknown_138 | 20.178 | 0 | 354.39 |
| 35 | ASTAN | CC_unknown_14 | 13.808 | 4E-04 | 363.25 |
| 36 | ASTAN | CC_unknown_141 | 11.711 | 0.0031 | 363.25 |
| 37 | ASTAN | CC_unknown_142 | 18.257 | 0 | 183.92 |
| 38 | ASTAN | CC_unknown_143 | 57.76 | 0 | 372.56 |
| 39 | ASTAN | CC_unknown_145 | 13.188 | 7E-04 | 363.25 |
| 40 | ASTAN | CC_unknown_146 | 12.62 | 0.0012 | 354.39 |
| 41 | ASTAN | CC_unknown_147 | 17.288 | 0 | 372.56 |
| 42 | ASTAN | CC_unknown_149 | 48.702 | 0 | 372.56 |
| 43 | ASTAN | CC_unknown_15 | 17.183 | 0 | 372.56 |
| 44 | ASTAN | CC_unknown_151 | 10.445 | 0.0108 | 363.25 |
| 45 | ASTAN | CC_unknown_153 | 15.689 | 1E-04 | 372.56 |
| 46 | ASTAN | CC_unknown_154 | 41.522 | 0 | 363.25 |
| 47 | ASTAN | CC_unknown_155 | 18.615 | 0 | 372.56 |
| 48 | ASTAN | CC_unknown_158 | 21.839 | 0 | 363.25 |
| 49 | ASTAN | CC_unknown_16 | 10.074 | 0.0156 | 1816.2 |
| 50 | ASTAN | CC_unknown_164 | 22.278 | 0 | 372.56 |
| 51 | ASTAN | CC_unknown_167 | 39.31 | 0 | 372.56 |
| 52 | ASTAN | CC_unknown_168 | 10.56 | 0.0097 | 392.7 |
| 53 | ASTAN | CC_unknown_169 | 10.356 | 0.0118 | 372.56 |
| 54 | ASTAN | CC_unknown_17 | 52.237 | 0 | 372.56 |
| 55 | ASTAN | CC_unknown_170 | 57.889 | 0 | 363.25 |
| 56 | ASTAN | CC_unknown_172 | 33.36 | 0 | 181.62 |
| 57 | ASTAN | CC_unknown_174 | 10.436 | 0.0109 | 354.39 |
| 58 | ASTAN | CC_unknown_181 | 18.714 | 0 | 372.56 |
| 59 | ASTAN | CC_unknown_182 | 11.327 | 0.0045 | 363.25 |
| 60 | ASTAN | CC_unknown_183 | 63.614 | 0 | 363.25 |
| 61 | ASTAN | CC_unknown_185 | 12.843 | 0.001 | 372.56 |
| 62 | ASTAN | CC_unknown_186 | 10.663 | 0.0087 | 363.25 |
| 63 | ASTAN | CC_unknown_189 | 73.299 | 0 | 363.25 |
| 64 | ASTAN | CC_unknown_19 | 10.776 | 0.0078 | 363.25 |
| 65 | ASTAN | CC_unknown_191 | 58.068 | 0 | 363.25 |
| 66 | ASTAN | CC_unknown_194 | 13.648 | 4E-04 | 363.25 |

| | | | | | |
|---|---|---|---|---|---|
| 67 | ASTAN | CC_unknown_199 | 84.242 | 0 | 363.25 |
| 68 | ASTAN | CC_unknown_2 | 21.007 | 0 | 2421.7 |
| 69 | ASTAN | CC_unknown_20 | 34.274 | 0 | 363.25 |
| 70 | ASTAN | CC_unknown_200 | 11.665 | 0.0032 | 363.25 |
| 71 | ASTAN | CC_unknown_203 | 67.627 | 0 | 363.25 |
| 72 | ASTAN | CC_unknown_204 | 10.212 | 0.0136 | 354.39 |
| 73 | ASTAN | CC_unknown_205 | 10.007 | 0.0167 | 354.39 |
| 74 | ASTAN | CC_unknown_21 | 11.216 | 0.005 | 363.25 |
| 75 | ASTAN | CC_unknown_215 | 10.581 | 0.0095 | 372.56 |
| 76 | ASTAN | CC_unknown_22 | 12.19 | 0.0019 | 372.56 |
| 77 | ASTAN | CC_unknown_227 | 13.81 | 4E-04 | 363.25 |
| 78 | ASTAN | CC_unknown_229 | 10.328 | 0.0121 | 363.25 |
| 79 | ASTAN | CC_unknown_231 | 12.65 | 0.0012 | 354.39 |
| 80 | ASTAN | CC_unknown_24 | 17.843 | 0 | 372.56 |
| 81 | ASTAN | CC_unknown_243 | 13.767 | 4E-04 | 2421.7 |
| 82 | ASTAN | CC_unknown_250 | 49.844 | 0 | 363.25 |
| 83 | ASTAN | CC_unknown_255 | 12.458 | 0.0014 | 181.62 |
| 84 | ASTAN | CC_unknown_257 | 77.824 | 0 | 363.25 |
| 85 | ASTAN | CC_unknown_26 | 39.7 | 0 | 354.39 |
| 86 | ASTAN | CC_unknown_260 | 13.466 | 5E-04 | 363.25 |
| 87 | ASTAN | CC_unknown_264 | 43.049 | 0 | 372.56 |
| 88 | ASTAN | CC_unknown_27 | 10.072 | 0.0156 | 16.06 |
| 89 | ASTAN | CC_unknown_272 | 73.939 | 0 | 372.56 |
| 90 | ASTAN | CC_unknown_28 | 12.63 | 0.0012 | 363.25 |
| 91 | ASTAN | CC_unknown_283 | 15.732 | 1E-04 | 2906 |
| 92 | ASTAN | CC_unknown_299 | 10.271 | 0.0128 | 363.25 |
| 93 | ASTAN | CC_unknown_3 | 17.043 | 0 | 363.25 |
| 94 | ASTAN | CC_unknown_30 | 17.471 | 0 | 1614.4 |
| 95 | ASTAN | CC_unknown_300 | 55.067 | 0 | 363.25 |
| 96 | ASTAN | CC_unknown_306 | 13.967 | 3E-04 | 354.39 |
| 97 | ASTAN | CC_unknown_31 | 12.441 | 0.0015 | 726.5 |
| 98 | ASTAN | CC_unknown_313 | 25.045 | 0 | 363.25 |
| 99 | ASTAN | CC_unknown_32 | 11.778 | 0.0029 | 363.25 |
| 100 | ASTAN | CC_unknown_325 | 45.552 | 0 | 354.39 |

**Table S5:** List of CCs selected both by Escouffier's equivalent vectors and LSP methods.

| | Dataset | CC_id | Escouffier | RV | PNmax | Pvalue | Period | Periodicity |
|---|---|---|---|---|---|---|---|---|
| **1** | ASTAN | CC_unknown_535 | 1 | 0.248 | 99.725 | 0 | 372.56 | 1 |
| **2** | ASTAN | CC_unknown_154 | 2 | 0.32 | 41.522 | 0 | 363.25 | 1 |
| **3** | ASTAN | CC_unknown_532 | 3 | 0.372 | 89.553 | 0 | 363.25 | 1 |
| **4** | ASTAN | CC_unknown_168 | 4 | 0.409 | 10.56 | 0.0097 | 392.7 | 1 |
| **5** | ASTAN | CC_unknown_547 | 5 | 0.442 | 13.203 | 7E-04 | 354.39 | 1 |
| **7** | ASTAN | CC_unknown_13 | 7 | 0.491 | 102.62 | 0 | 363.25 | 1 |
| **10** | ASTAN | CC_unknown_257 | 10 | 0.544 | 77.824 | 0 | 363.25 | 1 |
| **11** | ASTAN | CC_unknown_498 | 11 | 0.557 | 33.229 | 0 | 363.25 | 1 |
| **12** | ASTAN | CC_unknown_370 | 12 | 0.57 | 40.716 | 0 | 363.25 | 1 |
| **14** | ASTAN | CC_unknown_172 | 14 | 0.592 | 33.36 | 0 | 181.62 | 1 |
| **18** | ASTAN | CC_unknown_183 | 18 | 0.627 | 63.614 | 0 | 363.25 | 1 |
| **19** | ASTAN | CC_unknown_550 | 19 | 0.634 | 45.499 | 0 | 363.25 | 1 |
| **20** | ASTAN | CC_unknown_35 | 20 | 0.641 | 11.1 | 0.0056 | 2906 | 1 |
| **21** | ASTAN | CC_unknown_30 | 21 | 0.648 | 17.471 | 0 | 1614.4 | 1 |
| **23** | ASTAN | CC_unknown_80 | 23 | 0.661 | 16.021 | 0 | 363.25 | 1 |
| **26** | ASTAN | CC_unknown_462 | 26 | 0.678 | 19.424 | 0 | 354.39 | 1 |
| **27** | ASTAN | CC_unknown_530 | 27 | 0.683 | 32.635 | 0 | 354.39 | 1 |
| **28** | ASTAN | CC_unknown_126 | 28 | 0.689 | 30.339 | 0 | 2906 | 1 |
| **31** | ASTAN | CC_unknown_134 | 31 | 0.702 | 47.33 | 0 | 363.25 | 1 |
| **33** | ASTAN | CC_unknown_272 | 33 | 0.71 | 73.939 | 0 | 372.56 | 1 |
| **36** | ASTAN | CC_unknown_483 | 36 | 0.721 | 39.84 | 0 | 354.39 | 1 |
| **37** | ASTAN | CC_unknown_553 | 37 | 0.725 | 10.356 | 0.0118 | 16.06 | 1 |
| **38** | ASTAN | CC_unknown_20 | 38 | 0.728 | 34.274 | 0 | 363.25 | 1 |
| **39** | ASTAN | CC_unknown_12 | 39 | 0.731 | 13.212 | 7E-04 | 354.39 | 1 |
| **41** | ASTAN | CC_unknown_58 | 41 | 0.737 | 15.912 | 0 | 354.39 | 1 |
| **42** | ASTAN | CC_unknown_62 | 42 | 0.74 | 30.798 | 0 | 807.22 | 1 |
| **44** | ASTAN | CC_unknown_227 | 44 | 0.746 | 13.81 | 4E-04 | 363.25 | 1 |
| **48** | BBMO | CC_unknown_112 | 3 | 0.403 | 16.829 | 0 | 515.86 | 1 |
| **50** | BBMO | CC_unknown_213 | 5 | 0.478 | 11.852 | 0.0023 | 487.97 | 1 |
| **53** | BBMO | CC_unknown_401 | 8 | 0.552 | 14.165 | 2E-04 | 501.53 | 1 |
| **57** | BBMO | CC_unknown_389 | 12 | 0.604 | 10.489 | 0.0089 | 501.53 | 1 |
| **58** | BBMO | CC_unknown_154 | 13 | 0.615 | 15.424 | 1E-04 | 515.86 | 1 |

| | | | | | | | | |
|---|---|---|---|---|---|---|---|---|
| 79 | BBMO | CC_unknown_296 | 34 | 0.74 | 20.699 | 0 | 501.53 | 1 |
| 81 | BBMO | CC_unknown_126 | 36 | 0.747 | 13.3 | 5E-04 | 515.86 | 1 |
| 82 | SOLA | CC_unknown_425 | 1 | 0.341 | 16.862 | 0 | 360.5 | 1 |
| 85 | SOLA | CC_unknown_126 | 4 | 0.551 | 23.175 | 0 | 360.5 | 1 |
| 87 | SOLA | CC_unknown_149 | 6 | 0.604 | 14.401 | 1E-04 | 372.93 | 1 |
| 94 | SOLA | CC_unknown_143 | 13 | 0.707 | 12.967 | 3E-04 | 360.5 | 1 |
| 95 | SOLA | CC_unknown_183 | 14 | 0.717 | 16.434 | 0 | 360.5 | 1 |

**Table S6:** Summary information of temporal analysis.

| | ASTAN | BBMO | SOLA | Sum |
|---|---|---|---|---|
| **Nb Syndiniales CCs** | 572 | 2171 | 566 | 3309 |
| **Nb unassigned Syndiniales CCs** | 558 | 2142 | 556 | 3256 |
| **Nb CCs selected by Escoufier vectors** | 45 | 36 | 17 | 98 |
| **Nb CCs selected by Lomb-Scargle algorithm** | 208 | 118 | 15 | 341 |
| **Average recurrence period (days)** | 570 | 578 | 365 | - |
| **Min recurrence period (days)** | 16 | 23 | 360 | - |
| **Max recurrence period (days)** | 2906 | 3611 | 372 | - |
| **Nb CCs selected by Escoufiers vectors and Lomb-Scargle algorithm*** | 27 | 7 | 5 | 39 |
| **Nb selected* CCs present in the three time-series** | 1 | 1 | 1 | - |
| **Selected* CCs ids present in the three time-series** | CC_unknown_126 | CC_unknown_126 | CC_unknown_126 | - |
| **Recurrence period (days)** | 2906 | 515 | 360 | - |
| **Nb selected* CCs present in ASTAN and BBMO** | 1 | 1 | - | - |
| **Selected* CCs ids present in ASTAN and BBMO** | CC_unknown_154 | CC_unknown_154 | - | - |
| **Recurrence period (days)** | 363 | 515 | - | - |
| **Nb selected* CCs present in ASTAN and SOLA** | 1 | - | 1 | - |
| **Selected* CCs ids present in ASTAN and SOLA** | CC_unknown_183 | - | CC_unknown_183 | - |
| **Recurrence period (days)** | 360 | - | 363 | - |
| **Nb selected* CCs present in BBMO and SOLA** | - | 0 | 0 | - |
| **Selected* CCs ids present in BBMO and SOLA** | - | - | - | - |
| **Nb selected* CCs endemic to time-series** | 24 | 5 | 3 | 32 |

**Table S7:** Abundance and seasonality of CCs selected both Escouffier's equivalent vectors and LSP methods.

| | Dataset | CC_id | mean_abundance | Year | Season |
|---|---|---|---|---|---|
| 1 | ASTAN | CC_unknown_12 | 0.05 | 2009 | 1 |
| 2 | ASTAN | CC_unknown_12 | 0.05 | 2010 | 1 |
| 3 | ASTAN | CC_unknown_12 | 0.05 | 2011 | NA |
| 4 | ASTAN | CC_unknown_12 | 0.05 | 2012 | NA |
| 5 | ASTAN | CC_unknown_12 | 0.05 | 2013 | 1 |
| 6 | ASTAN | CC_unknown_12 | 0.05 | 2014 | 2 |
| 7 | ASTAN | CC_unknown_12 | 0.05 | 2015 | 1 |
| 8 | ASTAN | CC_unknown_12 | 0.05 | 2016 | 2 |
| 9 | ASTAN | CC_unknown_13 | 0.98 | 2009 | 3 |
| 10 | ASTAN | CC_unknown_13 | 0.98 | 2010 | 4 |
| 11 | ASTAN | CC_unknown_13 | 0.98 | 2011 | 4 |
| 12 | ASTAN | CC_unknown_13 | 0.98 | 2012 | 4 |
| 13 | ASTAN | CC_unknown_13 | 0.98 | 2013 | 4 |
| 14 | ASTAN | CC_unknown_13 | 0.98 | 2014 | 3 |
| 15 | ASTAN | CC_unknown_13 | 0.98 | 2015 | 4 |
| 16 | ASTAN | CC_unknown_13 | 0.98 | 2016 | 4 |
| 17 | ASTAN | CC_unknown_20 | 0.18 | 2009 | 3 |
| 18 | ASTAN | CC_unknown_20 | 0.18 | 2010 | 3 |
| 19 | ASTAN | CC_unknown_20 | 0.18 | 2011 | 3 |
| 20 | ASTAN | CC_unknown_20 | 0.18 | 2012 | 3 |
| 21 | ASTAN | CC_unknown_20 | 0.18 | 2013 | 2 |
| 22 | ASTAN | CC_unknown_20 | 0.18 | 2014 | 2 |
| 23 | ASTAN | CC_unknown_20 | 0.18 | 2015 | NA |
| 24 | ASTAN | CC_unknown_20 | 0.18 | 2016 | 2 |
| 25 | ASTAN | CC_unknown_30 | NA | 2009 | 3 |
| 26 | ASTAN | CC_unknown_30 | NA | 2010 | 1 |
| 27 | ASTAN | CC_unknown_30 | NA | 2011 | 1 |
| 28 | ASTAN | CC_unknown_30 | NA | 2012 | 1 |
| 29 | ASTAN | CC_unknown_30 | NA | 2013 | NA |
| 30 | ASTAN | CC_unknown_30 | NA | 2014 | NA |
| 31 | ASTAN | CC_unknown_30 | NA | 2015 | NA |
| 32 | ASTAN | CC_unknown_30 | NA | 2016 | NA |

| | | | | | |
|---|---|---|---|---|---|
| 33 | ASTAN | CC_unknown_35 | 0.03 | 2009 | 1 |
| 34 | ASTAN | CC_unknown_35 | 0.03 | 2010 | 4 |
| 35 | ASTAN | CC_unknown_35 | 0.03 | 2011 | 2 |
| 36 | ASTAN | CC_unknown_35 | 0.03 | 2012 | 2 |
| 37 | ASTAN | CC_unknown_35 | 0.03 | 2013 | 1 |
| 38 | ASTAN | CC_unknown_35 | 0.03 | 2014 | 2 |
| 39 | ASTAN | CC_unknown_35 | 0.03 | 2015 | 2 |
| 40 | ASTAN | CC_unknown_35 | 0.03 | 2016 | 1 |
| 41 | ASTAN | CC_unknown_58 | 0.01 | 2009 | 3 |
| 42 | ASTAN | CC_unknown_58 | 0.01 | 2010 | 2 |
| 43 | ASTAN | CC_unknown_58 | 0.01 | 2011 | 2 |
| 44 | ASTAN | CC_unknown_58 | 0.01 | 2012 | 1 |
| 45 | ASTAN | CC_unknown_58 | 0.01 | 2013 | 1 |
| 46 | ASTAN | CC_unknown_58 | 0.01 | 2014 | NA |
| 47 | ASTAN | CC_unknown_58 | 0.01 | 2015 | NA |
| 48 | ASTAN | CC_unknown_58 | 0.01 | 2016 | 1 |
| 49 | ASTAN | CC_unknown_62 | 0.06 | 2009 | 2 |
| 50 | ASTAN | CC_unknown_62 | 0.06 | 2010 | 3 |
| 51 | ASTAN | CC_unknown_62 | 0.06 | 2011 | 2 |
| 52 | ASTAN | CC_unknown_62 | 0.06 | 2012 | 4 |
| 53 | ASTAN | CC_unknown_62 | 0.06 | 2013 | 3 |
| 54 | ASTAN | CC_unknown_62 | 0.06 | 2014 | 3 |
| 55 | ASTAN | CC_unknown_62 | 0.06 | 2015 | 4 |
| 56 | ASTAN | CC_unknown_62 | 0.06 | 2016 | 2 |
| 57 | ASTAN | CC_unknown_80 | 0.06 | 2009 | 2 |
| 58 | ASTAN | CC_unknown_80 | 0.06 | 2010 | 2 |
| 59 | ASTAN | CC_unknown_80 | 0.06 | 2011 | 3 |
| 60 | ASTAN | CC_unknown_80 | 0.06 | 2012 | 3 |
| 61 | ASTAN | CC_unknown_80 | 0.06 | 2013 | 2 |
| 62 | ASTAN | CC_unknown_80 | 0.06 | 2014 | 2 |
| 63 | ASTAN | CC_unknown_80 | 0.06 | 2015 | 3 |
| 64 | ASTAN | CC_unknown_80 | 0.06 | 2016 | 4 |
| 65 | ASTAN | CC_unknown_126 | 0.94 | 2009 | 4 |
| 66 | ASTAN | CC_unknown_126 | 0.94 | 2010 | 4 |

| | | | | | |
|---|---|---|---|---|---|
| 67 | ASTAN | CC_unknown_126 | 0.94 | 2011 | 4 |
| 68 | ASTAN | CC_unknown_126 | 0.94 | 2012 | 4 |
| 69 | ASTAN | CC_unknown_126 | 0.94 | 2013 | 4 |
| 70 | ASTAN | CC_unknown_126 | 0.94 | 2014 | 4 |
| 71 | ASTAN | CC_unknown_126 | 0.94 | 2015 | 4 |
| 72 | ASTAN | CC_unknown_126 | 0.94 | 2016 | 4 |
| 73 | ASTAN | CC_unknown_134 | 0.39 | 2009 | 2 |
| 74 | ASTAN | CC_unknown_134 | 0.39 | 2010 | 4 |
| 75 | ASTAN | CC_unknown_134 | 0.39 | 2011 | 3 |
| 76 | ASTAN | CC_unknown_134 | 0.39 | 2012 | 2 |
| 77 | ASTAN | CC_unknown_134 | 0.39 | 2013 | 3 |
| 78 | ASTAN | CC_unknown_134 | 0.39 | 2014 | 4 |
| 79 | ASTAN | CC_unknown_134 | 0.39 | 2015 | 2 |
| 80 | ASTAN | CC_unknown_134 | 0.39 | 2016 | 2 |
| 81 | ASTAN | CC_unknown_154 | 0.28 | 2009 | NA |
| 82 | ASTAN | CC_unknown_154 | 0.28 | 2010 | 2 |
| 83 | ASTAN | CC_unknown_154 | 0.28 | 2011 | 2 |
| 84 | ASTAN | CC_unknown_154 | 0.28 | 2012 | 2 |
| 85 | ASTAN | CC_unknown_154 | 0.28 | 2013 | 2 |
| 86 | ASTAN | CC_unknown_154 | 0.28 | 2014 | 2 |
| 87 | ASTAN | CC_unknown_154 | 0.28 | 2015 | 2 |
| 88 | ASTAN | CC_unknown_154 | 0.28 | 2016 | 2 |
| 89 | ASTAN | CC_unknown_168 | NA | 2009 | 1 |
| 90 | ASTAN | CC_unknown_168 | NA | 2010 | 1 |
| 91 | ASTAN | CC_unknown_168 | NA | 2011 | NA |
| 92 | ASTAN | CC_unknown_168 | NA | 2012 | NA |
| 93 | ASTAN | CC_unknown_168 | NA | 2013 | NA |
| 94 | ASTAN | CC_unknown_168 | NA | 2014 | 2 |
| 95 | ASTAN | CC_unknown_168 | NA | 2015 | NA |
| 96 | ASTAN | CC_unknown_168 | NA | 2016 | 1 |
| 97 | ASTAN | CC_unknown_172 | 0.48 | 2009 | 3 |
| 98 | ASTAN | CC_unknown_172 | 0.48 | 2010 | 4 |
| 99 | ASTAN | CC_unknown_172 | 0.48 | 2011 | 4 |
| 100 | ASTAN | CC_unknown_172 | 0.48 | 2012 | 3 |
